# Representational learning of brain responses in executive function and higher-order cognition using deep graph convolutions

**DOI:** 10.1101/2021.07.26.453914

**Authors:** Yu Zhang, Nicolas Farrugia, Alain Dagher, Pierre Bellec

## Abstract

Brain decoding aims to infer human cognition from recordings of neural activity using modern neuroimaging techniques. Studies so far often concentrated on a limited number of cognitive states and aimed to classifying patterns of brain activity within a local area. This procedure demonstrated a great success on classifying motor and sensory processes but showed limited power over higher cognitive functions. In this work, we investigate a high-order graph convolution model, named ChebNet, to model the segregation and integration organizational principles in neural dynamics, and to decode brain activity across a large number of cognitive domains. By leveraging our prior knowledge on brain organization using a graph-based model, ChebNet graph convolution learns a new representation from task-evoked neural activity, which demonstrates a highly predictive signature of cognitive states and task performance. Our results reveal that between-network integration significantly boosts the decoding of high-order cognition such as visual working memory tasks, while the segregation of localized brain activity is sufficient to classify motor and sensory processes. Using twin and family data from the Human Connectome Project (n = 1,070), we provide evidence that individual variability in the graph representations of working-memory tasks are under genetic control and strongly associated with participants in-scanner behaviors. These findings uncover the essential role of functional integration in brain decoding, especially when decoding high-order cognition other than sensory and motor functions.

**Teaser:**

- Modelling functional integration through graph convolution is a necessary step towards decoding high-order human cognition.

**Significance statement:** Over the past two decades, many studies have applied multivariate pattern analysis to decode what task a human participant is performing, based on a scan of her brain. The vast majority of these studies have however concentrated on select regions and a specific domain, because of the computational complexity of handling full brain data in a multivariate model. With the fast progress in the field of deep learning, it is now possible to decode a variety of cognitive domains simultaneously using a full-brain model. By leveraging our prior knowledge on brain organization using a graph-based model, we uncovered different organizational principles in brain decoding for motor execution and high-order cognition by modelling functional integration through graph convolution.

## Introduction

Understanding the neural substrates of human cognition is a main goal of neuroscience research. Modern imaging techniques, such as functional magnetic resonance imaging (fMRI), provide an opportunity to map cognitive function in-vivo, and to decode the dynamics of cognitive processes from neural activity. Brain decoding has been an active topic since Haxby and colleagues first proposed the idea of using fMRI brain responses to predict the category of visual stimuli presented to a subject (1). Nowadays, a variety of computational models are used in the field, including multi-voxel pattern recognition, linear regression models, as well as nonlinear models such as deep artificial neural networks (DNN). Among which, DNN showed promising advantages over other linear models by providing an end-to-end solution to a direct mapping from recorded brain activity to brain cognition, for instance, using convolutional (2) and recurrent neural networks (3). However, most previous decoding studies aimed to segregate the spatial patterns of brain activation under different task conditions, but largely ignored the integration of brain dynamics during cognitive processes.

Functional segregation into highly localized brain areas, and functional integration at the levels of distributed brain regions, modules and networks, are fundamental principles of brain organization and have been widely observed in different populations and among a variety of cognitive tasks (4–7). So far, the majority of brain decoding studies only utilized the functional specialization hypothesis that aims to distinguish the localized brain activation patterns under a small number of experiment tasks, for instance, the involvement of the motor and sensory cortex during the movement of different body parts (8), or the engagement of different regions in the visual cortex for the recognition of various types of visual stimuli (9). As a result, such brain decoders were restricted to mostly motor and sensory processes (e.g. recognition of visual stimuli) and highly relied on domain knowledge (e.g. activating different parts of the visual cortex). This assumption of functional segregation also limited the generalizability of brain decoding towards high-order cognitive functions that were known to engage multiple brain systems. One typical example is the visual working memory task (VWM), for which multiple brain networks were involved through intense interactions among memory representations and other basic attention and sensory processes (10). For instance, early visual cortex played an important role in the detection of visual features including orientation (11), motion (12) and content (13), while the parietal and prefrontal cortex contributed to maintenance of visual information over a delayed interval (14). In these cases, both local and global information of brain activity may contribute to the decoding of cognitive processes (15,16).

We started to tackle this problem in our previous paper (17) by generalizing the convolutional operations from DNN onto brain organization. This approach can effectively capture both segregated brain activity of task-related brain regions, and information integration of neural dynamics within brain networks. Compared to previous linear and nonlinear decoding models, the proposed decoding model showed high generalizability over a variety of cognitive domains without relying on any prior information on the tested domain. However, this model relied on a simplified version of graph convolution which only took into account information integration within the same brain network at each layer. It showed limited power of representational learning on high-order cognitions that may involve complex forms of functional interactions across multiple brain systems.

To address this issue, we investigated a more sophisticated form of graph convolution in this study, namely ChebNet, which approximates the calculation of graph convolution using high-order Chebyshev polynomials. It has been proved that the ChebNet graph convolution is *K*-localized in space (on the graph) by taking up to *K*th order Chebychev polynomials (18). In other words, ChebNet integrates information within a relatively larger neighborhood by taking multiple steps of random walks on the brain graph. As a result, ChebNet graph convolution is capable of characterizing the complex forms of information processing during cognitive processes, i.e. segregating task-evoked activity from localized brain regions (K=0), integrating neural activity within the same brain networks (K=1), as well as information integration between different networks and among multiple brain systems (K>1). ChebNet provides a generalized form to encode this multiscale hierarchical organization of brain cognition in a single graph convolutional layer. The decoding model started with a parcellation that divides the whole brain into hundreds of brain regions and a brain graph that captures hierarchical and modular structures in brain organization. The brain graph as well as the dynamic information flow (i.e. task-evoked brain response at each brain region) on the graph was then mapped onto a new representational space through multilayer spatiotemporal graph convolutions.

These embedded graph representations naturally disassociate different cognitive tasks with large distances between task conditions and small distances within the same condition, and can improve the prediction of cognitive states by achieving better functional alignment between multiple trials and across different subjects.

In order to verify this hypothesis, we constructed the decoding model based on ChebNet graph convolution at different orders, ranging from local brain regions (K=0), to the same brain network (K=1), to multiple brain systems (K>1). All decoding models were evaluated on the task-fMRI database from the Human Connectome Project (HCP)(19) and simultaneously distinguished 6 cognitive domains or 21 task conditions by using a short time window of fMRI scans (e.g. 10 seconds). Under this framework, we systematically investigated how large-scale functional integration impact on brain decoding especially for multidomain decoding and decoding of high-order cognitive functions. Taking Motor and Working-memory tasks as examples, we further investigated the organizational structures among ChebNet layers within and across decoding models, and explored their relations to the two principles of brain organization, i.e. functional segregation and integration. Moreover, we investigated whether the representations learned through ChebNet graph convolutions were able to improve inter-subject alignment in brain responses and preserve individual variability in brain organization at the same time.

## Results

### Decoding cognitive functions with fine cognitive granularity and high accuracy

We proposed a decoding pipeline based on ChebNet graph convolution which automatically learns the spatiotemporal dynamics of brain activity from a short series of fMRI responses and predicts brain states based on learned feature representations (as shown in Figure 1-S1). The model starts with a brain graph with nodes representing brain parcels and edges representing brain connectivity, maps task-evoked fMRI responses onto the predefined brain graph, and learns high-level graph representations of neural activity by using stacked graph convolutions, taking into account both segregated neural activity within localized brain regions and functional interactions among between brain networks. For a more detailed description of the decoding model, please see the “Methods” section and Supplemental Information.

The ChebNet decoding model was evaluated using the cognitive battery of HCP task-fMRI dataset acquired from 1200 healthy subjects. Using the ChebNet-*K5* model (i.e. ChebNet graph convolution with K=5), the six cognitive domains were nearly perfectly differentiated from each other by only using 10s of brain recordings (approximately the shortest duration of task conditions in HCP), with an average test accuracy of 96% (mean=95.81%, STD =0.15% by using 10 fold cross-validation with shuffle splits). Moreover, the pipeline was capable of distinguishing experimental conditions with fine cognitive granularity and fine temporal resolution, either across multiple domains (see Table S3) or within each cognitive domain (see Table S2), and achieved high decoding accuracy on both tasks. Among the six cognitive domains (as shown in Figure 1-S2 and Table S2), the language tasks (2 conditions, story vs math), and motor tasks (5 conditions, left/right hand, left/right foot and tongue) were the most easily recognizable conditions, and showed the highest precision and recall scores (F1-score = 98.45% and 99.38%, respectively for classifying two language conditions and five motor conditions). The model achieved high decoding performance on other high-order cognitive functions when longer duration of task blocks was available, for instance working-memory (94.51%, classifying 8 conditions using 25s) and social cognition (96.58%, classifying 2 conditions using 23s). Our decoding model outperformed existing linear and nonlinear models including other deep learning architectures, which neglect the hierarchical brain organization during cognitive processes, for either classifying between cognitive domains (e.g. 93.7% when using 27 TRs reported in (2)) or decoding task states within specific domain (e.g. 92.6% and 92.9% for working-memory and social cognition respectively when decoding on 30s of fMRI data (3)).

### Brain decoding captured reliable and task-specific salient features

In order to validate that the decoding model used biological meaningful features, we generated the saliency maps on the trained decoding model by propagating the non-negative gradients backwards to the input layer (20). An input feature is *salient or important* only if its little variation causes big changes in the decoding output. The saliency scores were evaluated for each task trial independently and then averaged within each subject and for each condition (cognitive domain or task state). First, different sets of salient brain regions were detected for each cognitive domain (as shown in Figure 1C and D), for instance the involvement of the somatosensory cortex for motor execution (MOTOR) and the engagement of perisylvian language areas for language comprehension (LANGUAGE). Second, the salient features were not only highly selective to specific cognitive tasks but also very stable across trials and subjects. We took the Motor and Working-memory tasks as examples. The reliability of saliency values was evaluated by using repeated-measure ANOVA, controlling for the random effect of subjects and experimental trials. Only the salient brain regions that having high saliency values (>0.3) and showing a significant effect of task (*p* <0.001) were reported in the following analysis. As shown in Figure 2, salient brain regions in the sensorimotor cortex were identified in the Motor tasks, including region “a” (labelled as “area 5m” in the Glasser’s atlas) selectively activated during foot movements, region “b” (labelled as “area 2”) selectively activated during hand movements, regions c (labelled as “area OP4”) selectively activated during tongue movements. This distinctive pattern in the saliency maps were highly consistent across trials, sessions, and even subjects. For Working memory tasks, which involved both the differentiation between 0back vs 2back tasks and the recognition of different image categories, the decoding model learned reliable features related to both aspects, i.e. memory-load and image category. Here, we plotted the salient features for 0back and 2back tasks on face and place images. As shown in Figure 2D, ParaHippocampal Area 1 (PHA1) and ParaHippocampal Area 2 (PHA2) were selectively involved for the recognition of place images (repeated measure ANOVA, F-score=70.96 and 38.12, p-value=1.74e-8 and 3.0e-6 respectively for PHA1 and PHA2), while Fusiform Face Complex (FFC) and Lateral Occipital Area 1 (LO1) were selectively engaged for the recognition of faces (F-score=57.75 and 91.47, p-value=1.02e-7 and 1.75e-9 respectively for FFC and LO1). On other hand, for both place and face images, ParaHippocampal Area 3 (PHA3) was more involved in 0back tasks than 2back tasks (F-score=26.38, p-value=3.3e-5) while area PH was selectively engaged in the 2back tasks (F-score=102.56, p-value=6.01e-10) when fixing the stimuli category. Our results revealed that the decoding model captured reliable salient features from task-evoked brain activities, in order to distinguish among cognitive domains and task states. These salient features were derived from task-related brain regions and showed selective responses to different task conditions with high consistency not only with the same subject but also between different subjects (as shown in Figure 2), possibly revealing the biological basis of the decoding model.

**Figure 1.**
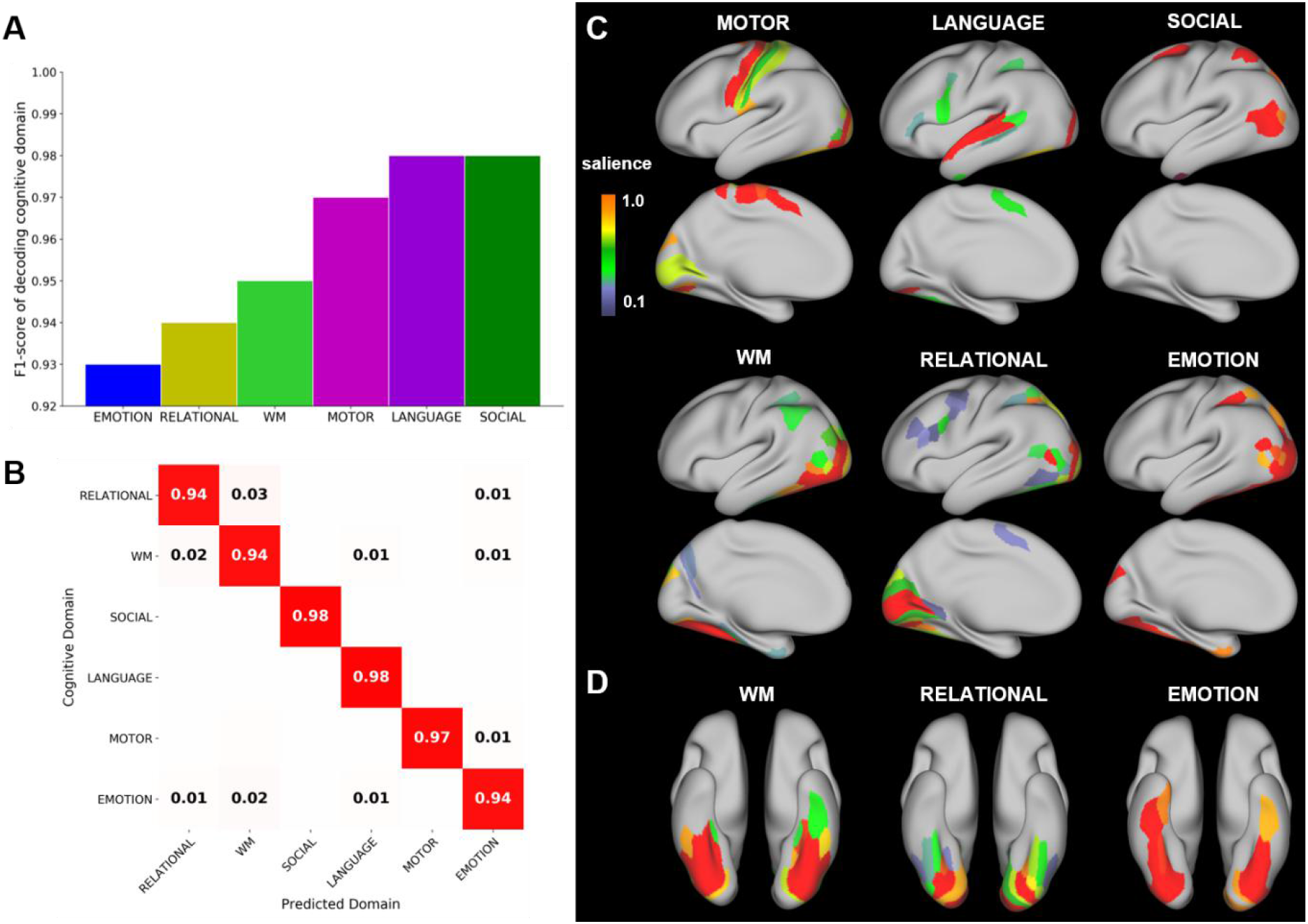
Decoding on six cognitive domains and the corresponding saliency maps. The decoding model predicted the cognitive domain from each 10s of fMRI responses and achieved an average test accuracy of 96%. The F1-score on each domain was shown in A. The corresponding cross-domain confusion matrix was shown in B. The saliency maps were evaluated for each task trial independently and then averaged within each subject and each domain. Different sets of salient brain regions were detected for each cognitive domain (C). Due to the similarity in task stimuli, the salient features in the ventral visual stream were identified for image recognition in three cognitive tasks, i.e. Working-memory (WM), relational processing (RELATIONAL) and emotional processing (EMOTION). Still, the decoding model captured different sets of visual areas for the three cognitive domains (D).

**Figure 2.**
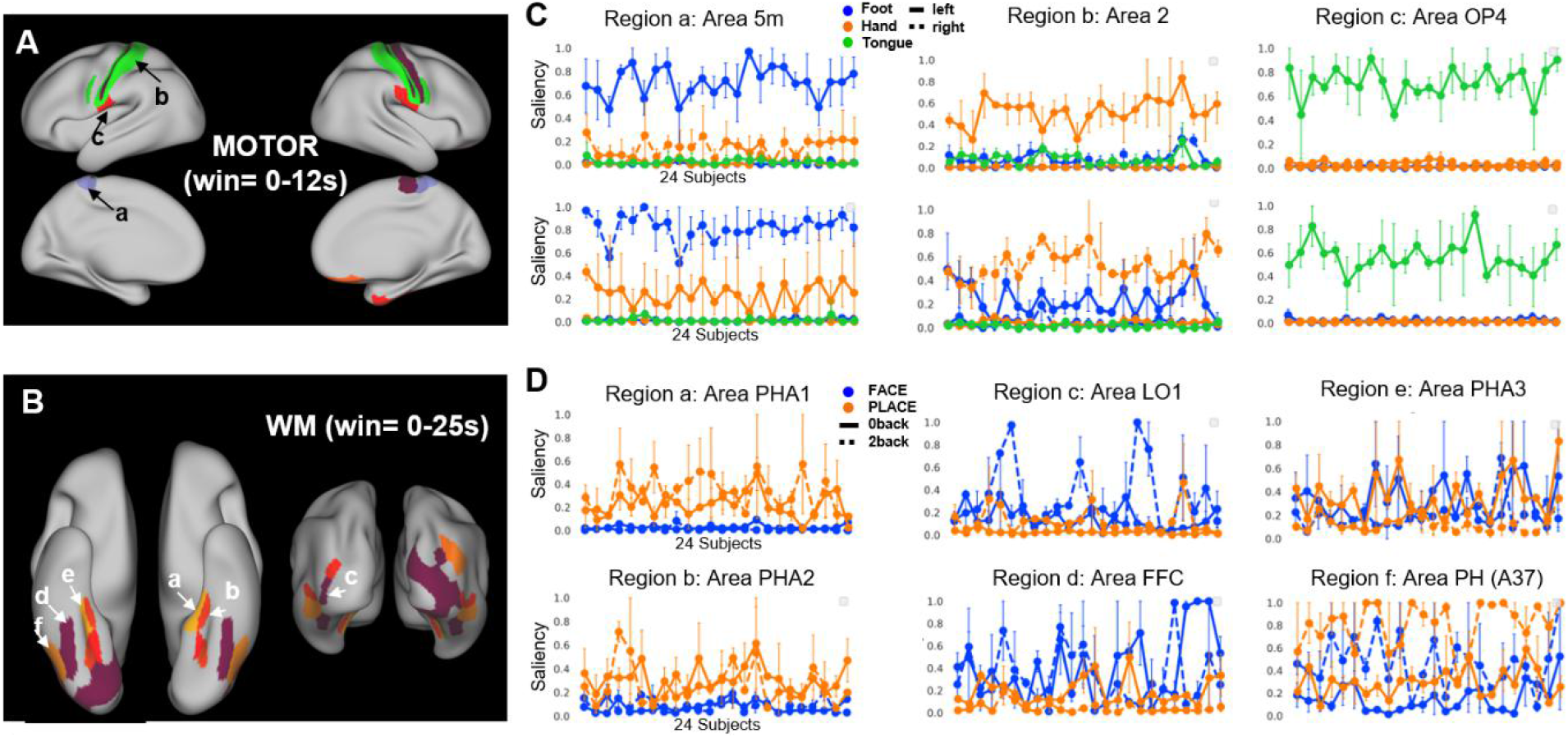
Salient features for the Motor (A) and Working-memory (B) tasks. Saliency value of each individual trial was estimated by using the guided backpropagation approach. The stability of saliency was evaluated by plotting the saliency values across randomly selected HCP subjects. The effect of task condition in the saliency values was then evaluated by using repeated-measure ANOVA, with the ‘subject’ as the random effect and ‘task condition’ as the within-subject effect. Only salient brain regions (saliency values>0.3, the full range of saliency is (0,1)) with a significant ‘task condition’ effect (p<0.001) were shown in the final saliency maps (A and B). For Motor task (C), three salient brain regions were selected that showed selective responses to the movement of foot (region “a”), hand (region “b”) and tongue (region “c”). The task trials corresponding to the movements of the left body parts were plotted in solid lines and in dashed lines for the right body parts. Brain regions in the left hemisphere were shown in the 1^st^ row and the right hemisphere shown in the 2^nd^ row. For Working-memory task (D), three sets of salient brain regions were selected that showed selective responses to the image category, e.g. place (1^st^ column, in orange) and face image (2^nd^ column, in blue), or to memory load, e.g. 0back (solid line) and 2back (3^rd^ column, dashed line).

### Decoding model learned hierarchical representations among ChebNet layers

The decoding model not only extracted biologically meaningful features associated with task-related brain regions, as illustrated by the saliency map analysis, but also learned hierarchical representations of brain response in each ChebNet layer. For instance, in the first graph convolutional layer (gcn1), the model learned various shapes of temporal kernels (accounting for the hemodynamic response in BOLD signals). Using these kernels, the model extracted a collection of spatial *“activation maps”*, which resembled the actual brain activation maps detected by the canonical GLM approach (Figure 3-S1 and Figure 3-S2). More sophisticated and task-specific feature representations were captured in deeper layers. In order to verify the hierarchy among ChebNet layers, we evaluated the similarity of feature representations using centered kernel alignment (CKA) with a linear kernel (21), with 0 < CKA < 1. As shown in Figure 3, a block-diagonal structure was detected in the CKA matrix of Working-memory (WM) tasks, indicating a hierarchical organization of representations among stacked ChebNet layers such that each layer inherited some information from previous layers, learned new representations in the current layer and passed these features onto the next layer.

**Figure 3.**
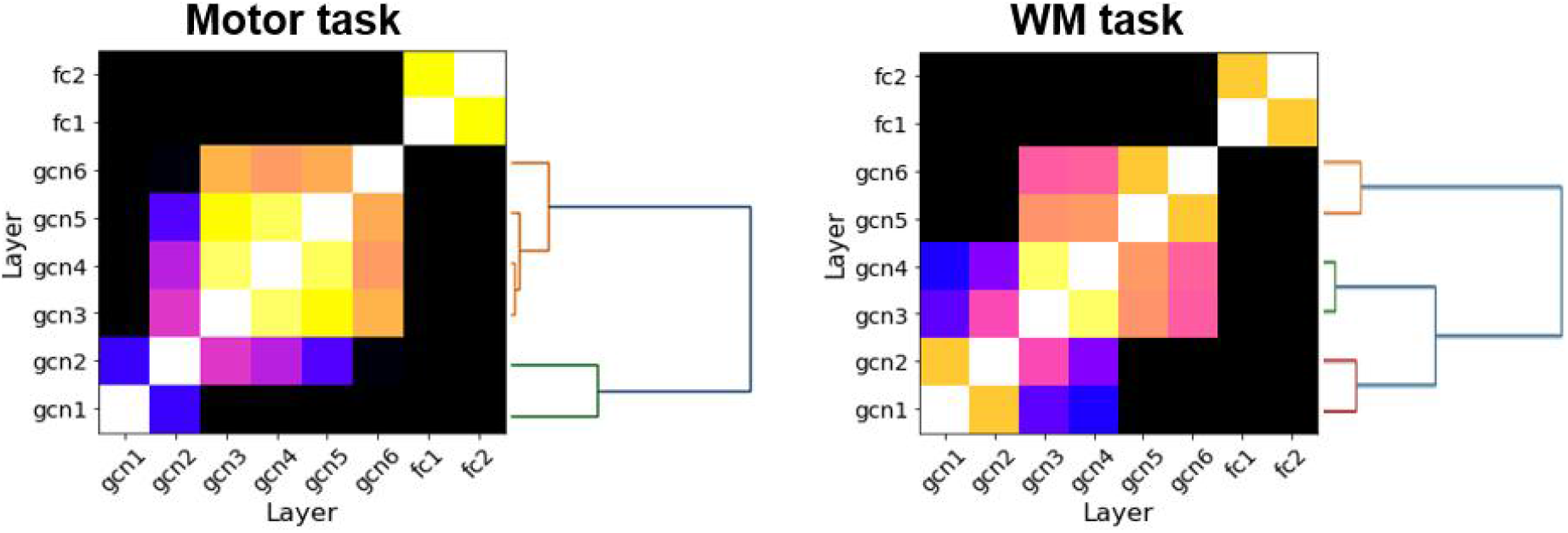
Hierarchical organization of layer representations learned through ChebNet graph convolutions. The similarity of representations between ChebNet layers was first calculated using CKA with a linear kernel. A distance metric was then generated from the CKA matrix (dis = 1-cka). After that, the hierarchical clustering was applied to the distance matrix using Ward’s linkage. The resulting dendrogram illustrated the hierarchical organization among ChebNet layers, for instance two-level organization in the Motor task and a tripartite organization in the Working-memory task.

The hierarchical clustering was applied to the CKA matrix and revealed a strong disassociation between the low-level features (gcn1 to gcn2), hidden representations (gcn3 to gcn4), and high-level representations (gcn5 to gcn6). Weak associations were detected across different levels (CKA=0.94 and 0.76 for within- and between-level similarity), with a stepwise progression towards the last ChebNet layer (CKA=0.54, 0.83, 0.92 for low, middle, high-level features as compared to gcn6), where category-specific information was present (i.e. different representations between task conditions). A similar hierarchical organizational structure was detected on the Motor task (as shown in Figure 3) but with fewer levels in the hierarchy, i.e. low- and high-level features, and with high redundancy in the middle layers (gcn3 to gcn5, average similarity with gcn6 is CKA=0.92). Still, distinct features were learned in the low- and high-level representations (CKA=0.58 for gcn1-gcn2 as compared to gcn6). Besides, the model already captured category-specific information starting in early ChebNet layers (Figure 6-S2). Our results indicated that the ChebNet decoding model learned hierarchical representations across graph convolutional layers in order to capture the underlying neural dynamics during cognitive processes. The hierarchy in the representations of the decoding model resembled the hierarchy in brain organization which has been reported in a variety of cognition functions especially for high-order cognition (22,23). Moreover, the different organizational patterns in ChebNet representations between cognitive tasks to some extent reflects different scales of information integration in cognitive processes, such that a deep ChebNet architecture was required to encode the complex forms of functional integration in WM, while a shallow ChebNet was sufficient to encode the segregation of localized brain activity during motor execution.

### Variable sensitivity to the K-order uncovers different organizational principles in cognitive processes

Another factor that impacts information integration in brain decoding is the *K*-order of ChebNet, by taking into account multi-level integration of neural dynamics at each graph convolutional layer, ranging from localized brain areas (K=0) to spatially distributed regions within the same network (K=1) and towards inter-connected brain networks (K>1). The choice of *K*-order not only showed an impact on the decoding performance, but also changed the hierarchy of feature representations learned in each ChebNet layer.

First of all, the decoding of six cognitive domains significantly impacted by the choice of *K*-order in ChebNet, indicating a faster convergence speed as well as higher decoding accuracy when using high-order models (Figure 4B). Significant improvements in decoding were detected between K=1 (integration of brain activity within the same network) and K>1 (between-network communication) (test accuracy = 93% vs 96% respectively for K=1 and K>1), significantly boosted compared to the localized decoding model (test accuracy = 83% for K=0). Second, variable sensitivity to the *K*-order was detected among different cognitive domains (Figure 4A). Specifically, for the Motor task, the decoding performance showed no improvement when increasing *K*, which means no gain from between-network communication during motor execution. Coinciding with this, the hierarchical organization of layer representations in the Motor task showed a very stable bipartition pattern when increasing K, i.e. low- and high-level features (as shown in Figure 4C). By contrast, the decoding of WM tasks gradually improved as increasing K and reaching the plateau after *K* > 5, which means that between-network communication and high-order integration plays an important role in WM, especially for distinguishing between 0back and 2back tasks (as shown in Figure 4-S1). Interestingly, the hierarchical organizational structure in WM (as shown in Figure 4D) started with three isolated clusters at K=1, gradually fused the representations by filling the gaps between neighboring layers, and converged to a stable tripartite organization at K=5 (i.e. low-, middle- and high-level representations). Further increase in the K-order did not change this organization but instead expanded the middle-level through encoding redundant hidden representations. Our results indicated that the variable sensitivity to the choice of *K*-order may uncover distinct organizational principles in cognitive processes, for instance, localized information processing within the motor and sensory cortex for motor execution, while complex forms of functional interaction and information integration across multiple brain systems/networks during WM tasks. Our findings coincided with the notion of functional segregation and integration in brain cognition (24), for instance, within-network communication is essential for motor execution, whereas integrative, between-network communication is critical for visual working memory (4).

**Figure 4.**
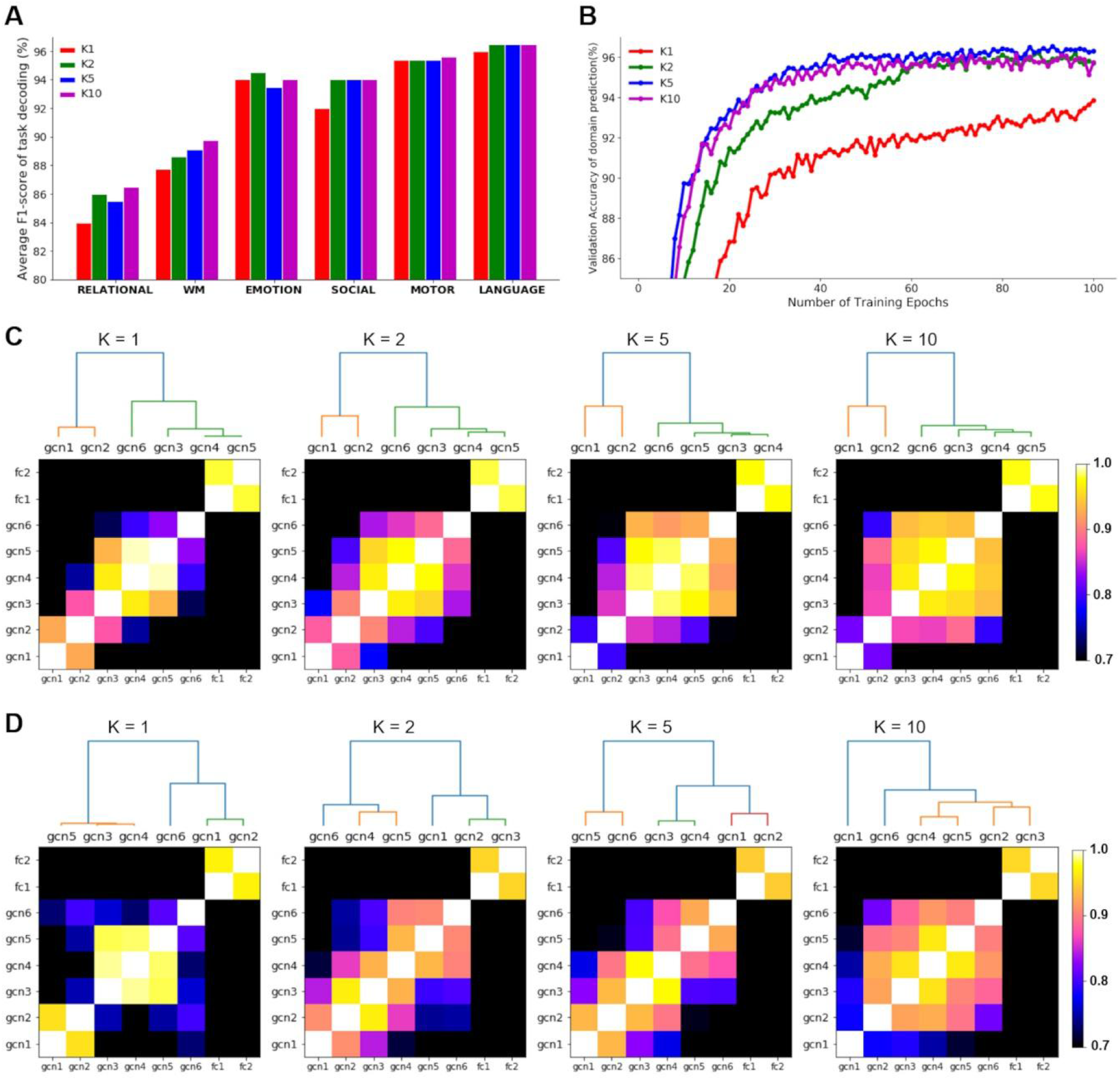
The effect of K-order on brain decoding and hierarchical organization of ChebNet. The effect of *K*-order on brain decoding was investigated by spanning over the list of [0,1,2,5,10]. The decoding performance on K=0 was not shown in this figure due to its low overall performance (decoding accuracy = 83.76%, 84.21%, 83.51% on training, validation and test sets). (B) High-order decoding models showed a faster convergence speed during model training and also achieved better decoding accuracy. Significant improvements were detected between K=1 (information integration within the same network) and K>1 (transmission of brain activity among inter-connected brain networks). (A) Variable sensitivity to the *K*-order was detected among different cognitive domains. The effect of *K*-order on each cognitive domain was estimated by averaging the F1-score on the test set. Among which, the Motor tasks showed stable decoding performance when increasing K while the decoding of WM tasks gradually improved as increasing *K*. (C) A stable two-level organization among ChebNet layers was revealed for the Motor tasks when increasing K. (D) For the Working-memory task, it started with an unstable bipartition and gradually evolved into a tripartite organization among ChebNet layers. The similarity of representations among ChebNet layers was calculated using CKA with a linear kernel. The hierarchical clustering was then applied to the distance matrix (dis = 1-cka) using Ward’s linkage and revealed the organizational principles among ChebNet layers.

### Functional integration in Working-memory tasks and segregation in Motor tasks

To further validate the functional segregation and integration hypothesis in brain decoding, we conducted a systematic analysis on the decoding models at different *K*-orders by calculating the similarity of representations between ChebNet models. We used the ChebNet-*K*5 model as the reference model for the similarity analysis.

For Motor tasks, the ChebNet-*K*1 model already captured the low-to-high-level organization in graph representations. Further increase in K did not change this organization but only caused redundant representations in the high-level features (average similarity with gcn6 in gcn3-gcn5 is CKA=0.78 and 0.92 for ChebNet-*K*1 and ChebNet-*K*5). A direct comparison between the two models (3rd row and 1st column in Figure 5B) revealed that, compared to the ChebNet-*K*5 model, the ChebNet-*K*1 model captured highly similar low-level features (CKA=0.92 for gcn1 when comparing between ChebNet-*K*1 and ChebNet-*K*5) and learned closely related high-level representations (CKA=0.84 for gcn6 between the two models). However, very different hidden representations were learned in the middle layers between the two models (averaged CKA=0.70 for gcn2 to gcn5). These results indicated that the highly segregated brain function, such as the sensory and motor tasks, did not involve high-level of information integration, but rather relied on neural transmission of brain activity within a local area or segregated networks.

**Figure 5.**
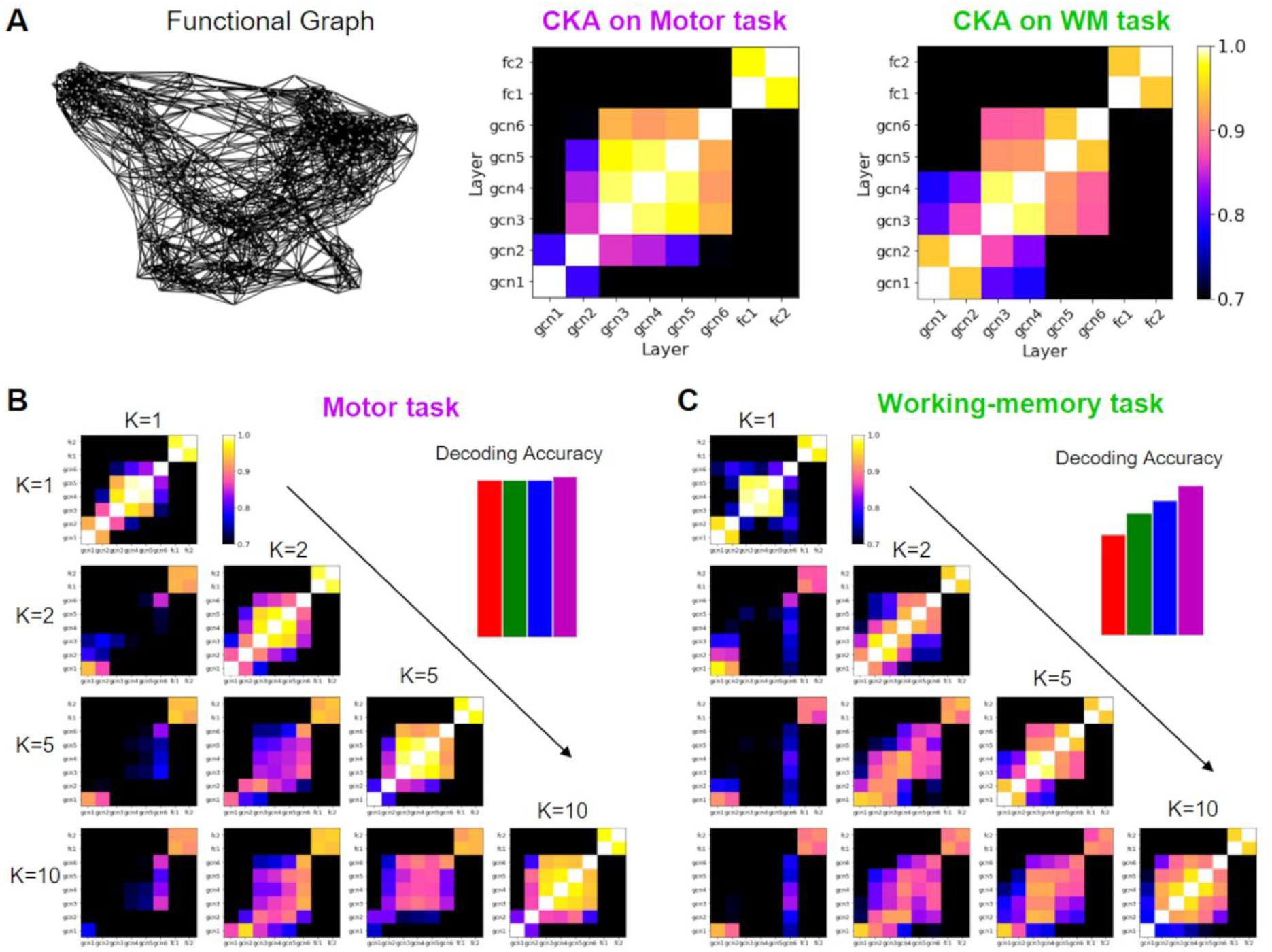
Similarity analysis of the decoding model with different *K* orders for the Motor and Working-memory tasks. The similarity analysis of layer representations demonstrated a hierarchical organization among stacked ChebNet layers in both Motor and Working-memory tasks (A). The decoding models were built on the same functional graph derived from the resting-state functional connectivity. The similarity of ChebNet representations was estimated not only between different layers in the same model but also between different models. For the Motor task (B), the decoding model already achieved the best performance at *K* = 1, and no further improvement on the decoding performance when increasing the *K*-order. In terms of layer representations, the ChebNet-*K*5 model captured similar low-level and high-level representations as ChebNet-*K*1 (3^rd^ row and 1^st^ column in B), but learned different representations in the hidden layers. Besides, higher redundancy was captured in the ChebNet-*K*5 model. This analysis indicated that ChebNet-*K*1 was enough to capture the functional segregation in brain activity during Motor tasks. For the Working-memory task (C), the decoding model showed high sensitivity to the choice of *K*-order and achieved the best decoding accuracy when *K* = 10. In terms of layer representations, the ChebNet-*K*1 model captured similar low-level representations as ChebNet-*K*5 (3^rd^ row and 1^st^ column in C), but learned very different hidden and high-level representations. On the other hand, ChebNet-*K*10 was highly similar to ChebNet-*K*5 (4^th^ row and 3^rd^ column in C), not only in the low-level representations (gcn1-gcn2), hidden representations (gcn3-gcn5), as well as high-level representations (gcn6). This analysis indicated that a high-order model was required in order to capture the complex forms of functional integration during Working-memory tasks.

On the other hand, for the Working Memory tasks, the ChebNet-*K*5 model captured a nice disassociation between low-level features (gcn1 to gcn2), hidden representations (gcn3 to gcn4), and high-level features (gcn5 to gcn6). Such hierarchical organization was broken in the ChebNet-*K*1 model due to poor between-layer communication (i.e. big gaps in the representations between neighboring layers, CKA=0.98 and 0.69 for within- and between-level similarity in ChebNet-*K*1). Moreover, the ChebNet-*K*1 model successfully captured the low-level features by showing high similarity to ChebNet-*K*5 in the first two ChebNet layers, but it was not capable of encoding high-level representations in the last ChebNet layer (3rd row and 1st column in Figure 5C, compared between ChebNet-*K*1 and ChebNet-*K*5, CKA=0.93 for gcn1, 0.88 for gcn2, 0.74 for gcn6). By contrast, the ChebNet-*K*10 model learned very similar representations in the low, middle and high ChebNet layers as in ChebNet-K5 (4th row and 3rd column in Figure 5C, compared between ChebNet-*K*5 and ChebNet-*K*10, CKA=0.94 for gcn1, 0.90 for gcn6, average CKA=0.90 for gcn3 to gcn5). These results indicated that the high-order cognitive functions required a large scale of information propagation and integration on the brain graph, not only involving the local connections within a specific brain network (*K* = 1) but also engaging the long-range connections across multiple networks (*K* ≥ 5).

### Improved functional alignment of cognitive states using graph convolution

The layer representations in ChebNet improved inter-subject alignment of task-evoked brain responses. For visualization purposes, we projected the feature representations of each ChebNet layer onto a 2-dimensional space using t-SNE (25). Compared to raw fMRI data or the activation maps derived from GLM analysis, the ChebNet representations of different task conditions were highly clustered and easily separated from each other, demonstrating a strong effect of task segregation (Figure 6A). The projections using other dimension reduction techniques were shown in Figure 6-S1, including PCA, UMAP (McInnes et al., 2018), and PHATE (Moon et al., 2019). The segregation effect was evaluated by calculating the modularity score (Q) of the state-transition graph on the projections of layer representations. As we went deeper along ChebNet layers, the segregation effect gradually strengthened and reached the peak in the last ChebNet layer. As shown in Figure 6-S2, for the Motor task, a low segregation was detected in the raw fMRI data (Q = 0.25), with slightly higher values in early ChebNet layers (e. g. Q = 0.41 in gcn1) and reaching the peak in the last ChebNet layer (Q = 0.60 in gcn6). A similar level of task segregation was observed when using a high-order ChebNet model, except for a faster convergence speed among ChebNet layers (Figure 6-S2B). For the WM tasks, the segregation effect was evaluated separately for the memory-load and image category. Interestingly, stronger segregation effect was detected among different image categories, e.g. place, face, body and tool images, than between different levels of memory loads, e.g. 0back and 2back (e.g. Q = 0.55 vs 0.38 in gcn6 respectively for the image category and memory load), but both higher than the effects in the raw fMRI data (Q = 0.06 vs 0.03 respectively).

**Figure 6.**
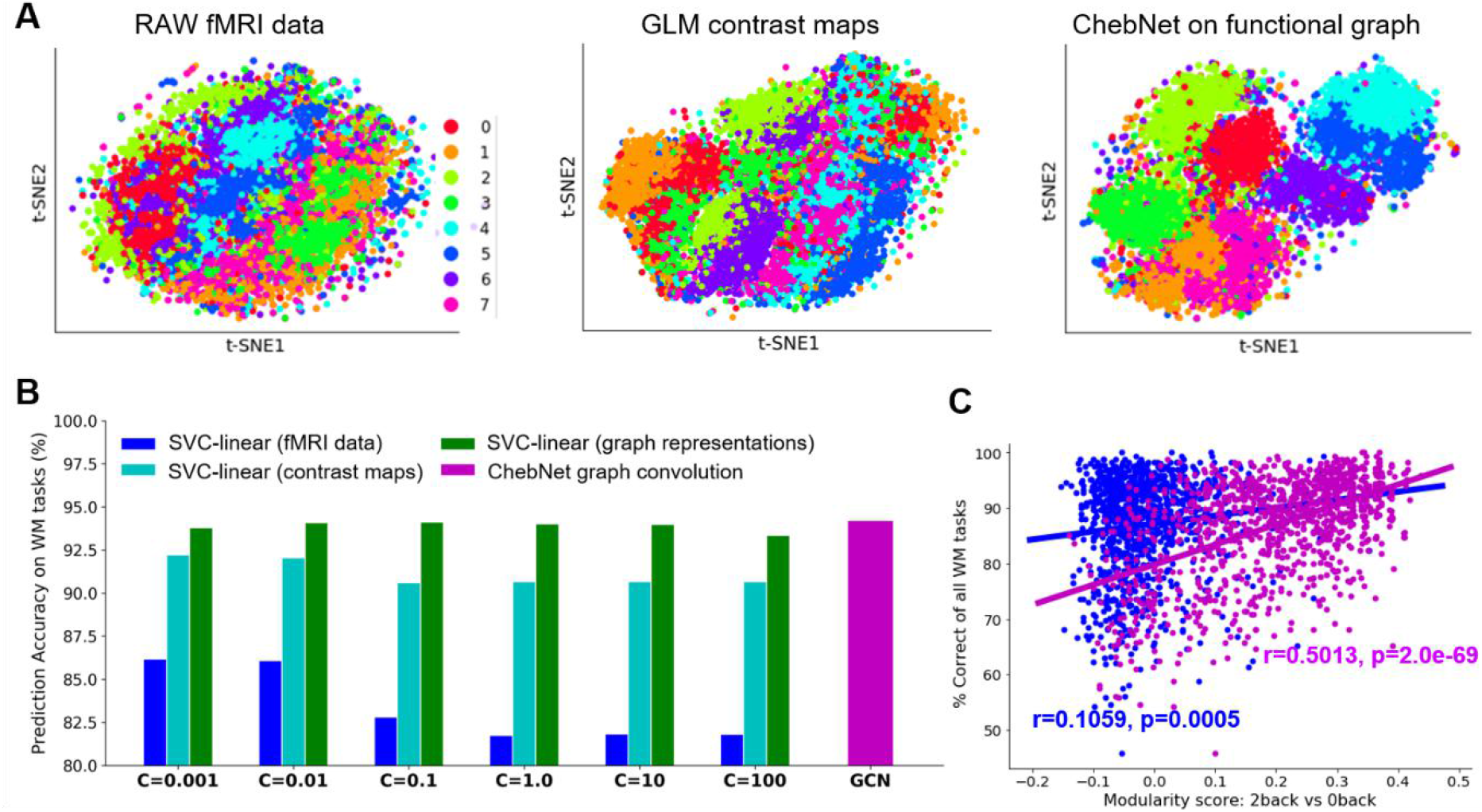
ChebNet graph representation improved the functional alignment of cognitive states, and induced higher decoding accuracy (B) and better prediction of task performance (C). Graph representations were extracted from the last ChebNet layer of the decoding model. Both fMRI data and contrast maps were mapped onto the same brain atlas, i.e. Glasser’s atlas (27) in the example, by averaging the fMRI time-series or z-scores of task activation within each brain region. (A) All three types of features, i.e. fMRI data, contrast maps and graph representations, were projected onto a 2-dimensional space using t-SNE (25) for the visualization purpose. Among them, graph representations showed high distinctions among different task conditions. (B) The decoding of eight working-memory tasks was re-evaluated by using multi-class support vector machine classification (SVC) on these features. Among them, graph representations showed the highest decoding accuracy, regardless of the chosen classifiers and parameters, e.g. linear classifier like SVC or nonlinear classifier such as ChebNet. (C) The effect of task segregation was evaluated by the modularity score on individual state-transition graph, as proposed by (26). We found a strong association between the task segregation of ChebNet representations and participants’ in-scanner task performance. The purple line indicated the association of participants’ in-scanner task performances with graph representations derived from ChebNet, while the blue line indicated the association with raw fMRI data.

### Association between ChebNet representations and behavioral performance

The segregation effect in the ChebNet representations not only boosted the decoding of cognitive stats, through better alignment of brain response across trials and subjects, but also improved the association between behavior and brain organization, largely preserving individual variability in cognitive processes. Specifically, the decoding model achieved much higher decoding accuracy when using the ChebNet representations as features, regardless of the choices of classifiers and parameters, for instance either using a multi-class support vector machine classification (SVC) or deep neural networks (as shown in Figure 6B). In addition, the segregation in the representations of brain response was significantly associated with behavioral performance during visual working memory task (as shown in Figure 6C). It has been shown in previous studies that the modularity of state-transition on individual fMRI data was significantly associated with participants’ in-scanner task performance (26). This association was also observed in our analysis. Moreover, we found a much stronger brain-behavior association when constructing the state-transition graph based on ChebNet representations rather than using raw fMRI data (Figure 6C). Specifically, the segregation of memory load in graph representations was highly associated with subjects’ in-scanner performance (as shown in Figure 7 and Figure 7-S1), including the average accuracy on all WM tasks (*r* =0.5013, *p* =2.0e-69), on 0back tasks (*r* =0.4450, *p* =2.33e-53) and on 2back tasks (*r* =0.3962, *p* =1.08e-13), as well as the reaction time on all WM tasks (*r* =-0.2601, *p* =5.74e-18), on 0back tasks (*r* =-0.3592, *p* =1.13e-33) and on 2back tasks (*r* =-0.1173, *p* = 0.0001). This analysis was done by using all subjects from the *HCP S1200* database (*N* =1074 of all subjects with available behavioral and imaging data for WM tasks). The significant correlations were sustained after controlling for the effect of confounds including age, gender, handedness and head motion (*r* =0.4659, *p* =5.74e-59 for the average accuracy; *r* =-0.2552, *p* =2.0e-16 for the reaction time). Moreover, the segregation of ChebNet representations as well as in-scanner behavioral performance during WM tasks were significantly heritable in HCP population (h^2^ =0.2882 for ChebNet representations, h^2^ =0.5624 and 0.4118 for average accuracy and reaction time in WM tasks, see Table S4 for all heritability estimates) and demonstrated significant shared genetic variance in ChebNet representations and behavioral scores (ρ_g_ = 0.80 and -0.39 respectively for the average accuracy and reaction time, see Table 1 for shared genetic influences in brain-behavioral associations).

**Figure 7.**
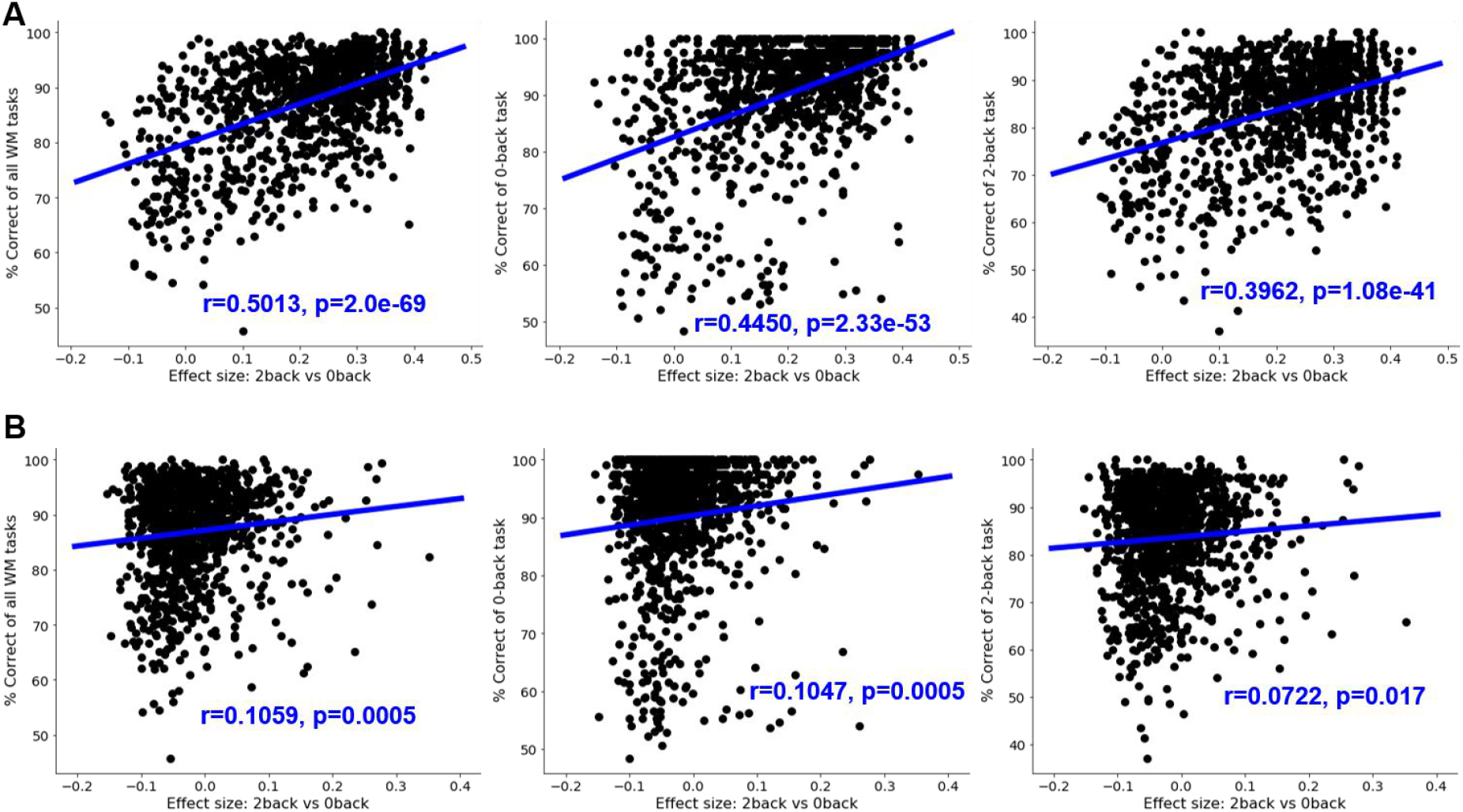
Modularity scores in the state-transition graph significantly correlated with correct responses during Working-memory tasks. The modularity score was calculated based on the state-transition graph of each subject, as proposed by (26). Specifically, we first constructed a kNN graph from the t-SNE projections of fMRI signals (B) or learned graph representations (A) of each subject. The modularity scores were then evaluated based on the kNN graph with the partition provided by task conditions (e.g. 0back vs 2back). We found significant correlations between the modularity scores of graph representations (A) and correct responses during working-memory tasks (1^st^ panel), 0back task conditions (2^nd^ panel) and 2back task conditions (3^rd^ panel). Much weaker associations were detected in the raw fMRI data (B). The blue lines indicated the linear regression models between the modularity score of the state-transition graph and the average accuracy during task performance. The analysis was done among all subjects from HCP S1200 release, with complete records of behavioral and imaging data for working memory tasks (N=1074).

**Table 1:** Shared genetic influences in ChebNet representations and behavioral scores. Bivariate genetic analyses were applied to quantify the shared genetic variance between ChebNet representations of brain responses and behavioral measures. Strong associations of ChebNet representations with the average accuracy (Acc) and reaction time (RT) during WM tasks were observed, mainly due to shared genetic effects in brain response and behaviors. Both genetic and phenotypic correlations reached a high-level of significance (FDR corrected). ***: *p* <0.001, **: *p* <0.01, *: p<0.05

|  | **Phenotypic**<br>**correlation ($\rho_p$)** | **Genetic**<br>**correlation ($\rho_g$)** |
| --- | --- | --- |
| WM_Task_Acc | 0.4659 *** | 0.7992 *** |
| WM_Task_2bk_Acc | 0.3716 *** | 0.7731 *** |
| WM_Task_0bk_Acc | 0.4189 *** | 0.8650 *** |
| WM_Task_RT | -0.2552 *** | -0.3895 ** |
| WM_Task_2bk_RT | -0.1173 *** | -0.2455 |
| WM_Task_0bk_RT | -0.3408 *** | -0.4967 ** |

## Discussion

We proposed a generalized framework for brain decoding based on ChebNet graph convolutions. The model takes in a short window of fMRI time series and a brain graph (with nodes representing brain parcels and edges representing brain connectivity), and then annotates brain activity with fine temporal resolution, and fine cognitive granularity. Using a 10s window of fMRI signals, our model identified 6 cognitive domains with a test accuracy of 96%, and distinguished fine-grained cognitive states on a trial basis with an accuracy above 93%, outperforming existing linear and nonlinear decoding models. This gain in brain decoding was mainly contributed by high-level integration of brain dynamics not only within limited subsets of brain regions but also between multiple brain networks. Specifically, we used high-order ChebNet graph convolution to encode the complex forms of functional integration during cognitive processes and captured hierarchical representations of brain activities at different levels. This hierarchical organizational pattern as well as the decoding accuracy was selectively impacted by the K-order in graph convolution, due to different organizational principles of the cognitive tasks. For segregated brain function like motor execution, the K=1 model achieved the best performance and revealed a stable 2-level hierarchy in neural representations. By contrast, for high-order cognition such as visual working-memory tasks, the model plateaued at K=5 and uncovered a tripartite organization in neural activity. Our findings revealed the essential role of functional integration in brain decoding, especially when decoding high-order cognition other than sensory and motor functions.

### Functional segregation and integration in brain decoding

Brain decoding has been a popular topic in neuroscience literature for decades since Haxby first proposed the idea of recognition of different visual stimuli using brain activity from the visual cortex (1,28). In the last decades, a variety of decoding models have been proposed with the aim to learn a linear discriminative function on the spatial patterns of brain activations associated with different task conditions. For instance, researchers have successfully attempted to use brain activity to reconstruct the frames of movies (29), or to decode the semantic context from words (30) and visual scenes (31) by using linear regression models. Recently, the fast development of deep artificial neural networks (DNNs) has also drawn a lot of attention in neuroscience research. Several different DNN architectures have been proposed to map human cognition from recorded brain activity, for instance using classical convolutional (2) and recurrent neural networks (3), or a generalized form of convolutions in the graph domain (17). However, the majority of brain decoding studies only utilized the functional specialization hypothesis that aims to distinguish the localized brain activation patterns by either training a linear classifier (16,32) or a nonlinear model through DNNs (2).

However, the majority of brain decoding studies so far only utilized the functional specialization hypothesis that aims to distinguish the localized brain activities from a single brain region or a small set of areas. This approach has shown promising results in the recognition of visual stimuli (28) and decoding the direction of finger movements (33). It may suffer from limited decoding power when dealing with large populations and a variety of cognitive states that involve not only motor and sensory perception but also high-order cognition (16). Such large-scale decoding is still challenging and may require a large collection of brain imaging data and incorporating brain responses from the whole brain in the decoding model, including both local and global information (34). So far, the functional integration at the whole-brain has been largely ignored in the brain decoding literature, but started to draw the attention of neuroscientists. Cole and colleagues (35) first showed that the information flow within functional networks was able to predict brain activation during cognitive tasks, specifically to predict activation patterns of unseen brain regions from regions of the same network. A similar idea was recently used in (3) by first extracting an integrated signal from each of 90 resting-state networks and then inferring brain states based on the temporal dependencies of these brain signals. Following this line of work, we recently proposed a graph convolutional network to decode brain states by propagating temporal dynamics of brain activity based on functional networks (17). In the present study we further extended this framework by exploring more variants in the graph convolutional network architecture. By using high-order graph convolutions, the model projected the spatiotemporal dynamics of cognitive processes onto a new representational space and integrated the context of brain activity in both local and global extent, ranging from brain region to functional networks and towards the whole brain.

Compared to previous linear and nonlinear decoding models, our proposed decoding model provided a generalized solution over a large population and a variety of cognitive domains. Besides, our model outperformed other approaches on the same dataset (as shown in Table S3), most of which followed the functional segregation assumption by predicting cognitive states from localized features of each brain parcel covering the entire cerebral cortex. After incorporating the network architecture of the human brain and integrating information flow within functional networks (e.g. first-order GCN), the classification accuracy was largely improved (90%, as stated in (17)). The decoding accuracy was further improved after taking into account the high-order interactions on the graph, not only within functional networks but also across multiple brain systems (93% using the 5-order ChebNet model).

Our results suggest that not only the segregated brain activations played an important part in distinguishing between cognitive processes as illustrated in previous brain decoding literature, but the functional integration within and between brain networks can also contribute to the classification of cognitive states to some degree. The tradeoff between functional segregation and integration largely depends on the nature of cognitive processes, for instance, localized brain signatures from motor and sensory cortex for Motor tasks (as shown in Figure 2A), while complex forms of functional interactions among multiple brain systems during WM tasks (as shown in Figure 2B). Their relations were automatically captured during the training of deep neural networks. Coinciding with this hypothesis, the decoding model achieved excellent performance in distinguishing different types of body movements when only considered the local context of brain activity either from a local area (94.7% in (2)) or within a functional network (96.6% in (3)). A similar level of performance was achieved when using either first-order or high-order graph convolutions (95.6% when using 10s fMRI signals). By contrast, when classifying 0-back and 2-back WM tasks, much higher classification errors were detected only using local brain activity (14% in (2)) compared to functional integration within the functional networks (10% in (3)). The classification errors among WM tasks were highly reduced when applying graph convolutions (<=9% when using ChebNet-K1, <=4% when using the ChebNet-K5, as shown in Figure 4-S1).

To conclude, our results demonstrated that an efficient brain decoder not only involved functional segregation, e.g. distinguishing localized brain activation during body movements, but also engaged functional integration, e.g. integrating brain activity among multiple brain networks during visual working memory tasks.

### Saliency of brain decoding goes beyond brain activation

Both saliency maps and brain activations aimed to reveal the neural substrates of cognitive processes. They also shared some common features, for instance, both relying on task-evoked brain responses and showing selective responses to different task conditions. However, their relations need to be addressed with caution. Brain activation was commonly used in neuroscience research to study the neural basis of cognitive processes by convolving neural activity with a canonical hemodynamic response function and to find the localization of each cognitive function using a generalized linear model (GLM) approach. However, as stated in Poldark’s paper (34), not all brain activations were “diagnostic” in terms of brain state prediction, i.e. distinguishing among different cognitive processes. To address this issue, we used saliency maps to detect the important features that show high contributions to the prediction.

First, common brain regions were revealed in both approaches, i.e. areas that not only strongly activated during task performance (in GLM analysis) but also largely contributed to the classification of different tasks (in saliency maps). For example, salient features in the sensorimotor cortex were detected for motor execution and in the ventral visual stream for image recognition during WM tasks (Figure 2 A and B). These brain regions have been well validated in the literature, that the primary motor cortex was engaged during movements of the human body (36) and ventral temporal cortex was responsible for the recognition of face and place images (37). Most previous decoding studies were based on this set of brain regions, for instance, using brain activity of the visual cortex to decode the category of seen images including faces vs objects as well as different animal species (28).

Second, some inconsistent findings were reported by the two techniques. On one hand, low saliency values did not mean no brain activation. Instead, the regions might be activated for all task conditions (i.e. low saliency but showing high activation on all tasks). For example, the visual cortex was consistently activated during the Motor task (38), in response to the presentation of the cue images and visual instructions (e.g. images of hand, foot and tongue in the cue phase). But this activation pattern was not related to actual movements and not informative to distinguish between hand and tongue movements, i.e. much lower decoding accuracy in the cue phase compared to the movement blocks, as indicated in the Figure 6A in (17). Consequently, the visual cortex was not detected in the saliency maps of Motor tasks (Figure 2A). On the other hand, some brain regions with absence of strong activations might show high saliency to the prediction (i.e. high saliency but with low activation scores). For instance, high saliency values were detected in the bilateral area OP4 for tongue movements and in the left area PHA3 for recognition of both place and face images. These patterns were not detected by random but instead highly consistent across a number of subjects (Figure 2C and D). By contrast, when using the GLM approach, these regions were not detected in either the group activation maps from HCP database (Figure 5-S1 in (17)) or meta-analysis from a collection of previous studies (Figure 5-S2 in (17)). Specifically, weak activation was detected in the left but not right OP4 for the tongue movement (T-score=4.6 and -1.66 respectively for the left and right OP4 in the contrast maps of tongue vs rest, using group activation maps from HCPS500 release), while area PHA3 was not even activated for the recognition of face and place images (T-score= -1.78 and -7.02 respectively for the contrast maps of face and place images vs rest). One possible explanation of this is that task-evoked activities in these brain regions did not follow the shape of the canonical hemodynamic response function and thus were not detected by the GLM approach and not shown in the contrast maps. However, these brain regions still showed distinctive patterns of response to different cognitive tasks, and thus were detected in the saliency maps when such constraint of temporal dynamics was not applied in the decoding model. Our results suggest that not only brain activations but also deactivations or even brain responses not shown in the contrast maps might highly contribute to the classification of cognitive states. In addition, the detected salient features were highly consistent across subjects and showed selective responses to different cognitive states, may uncover the biological basis of the decoding model and shed a light on the anatomical and functional substrates of cognitive processes.

To conclude, our results suggested that a more generalized framework was required for brain decoding which does not rely on localized brain activations or spatial patterns of contrast maps, but instead decoding information from task-evoked brain responses of the whole brain. Consequently, the model was able to distinguish the neural dynamics of various cognitive functions in both spatial and temporal domains and learn new representations of brain organization during cognitive processes (39). The spatiotemporal graph convolution provided a promising solution for this problem by leveraging our prior knowledge on brain organization using a graph-based model. Instead of classifying patterns of brain activity within a local area, graph convolution takes into account the functional interactions of neural dynamics across multiple networks and projects the spatiotemporal dynamics of cognitive processes onto a new representational space. Moreover, ChebNet graph convolution naturally incorporates both functional segregation and integration for brain decoding, i.e. distinguishing localized brain activities from a subset of brain regions or within a specific brain network (first-order convolution) and encoding complex forms of functional interactions among multiple brain systems (high-order convolution), in line with different organizational principles of cognitive processes. As a result, the learned representations improved the functional alignment among trials and subjects and therefore increased the decoding of cognitive states. More importantly, by largely preserving the individual variability in brain organization, the ChebNet representations achieved better associations with human behaviors during task, demonstrating shared genetic influences in brain responses and behaviors. The present work suggests the feasibility of large-scale multi-domain decoding with full-brain models, opening new avenues for modelling of naturalistic tasks such as movies or video games.

## Materials and Methods

### fMRI Datasets and Preprocessing

We used the block-design task-fMRI dataset from the Human Connectome Project S1200 release (https://db.humanconnectome.org/data/projects/HCP_1200). The minimal preprocessed fMRI data in CIFTI formats were selected. The preprocessing pipelines includes two steps (40): 1) fMRIVolume pipeline generates “minimally preprocessed” 4D time-series (i.e. “.nii.gz” file) that includes gradient unwarping, motion correction, fieldmap-based EPI distortion correction, brain-boundary-based registration of EPI to structural T1-weighted scan, non-linear (FNIRT) registration into MNI152 space, and grand-mean intensity normalization. 2) fMRISurface pipeline projects fMRI data from the cortical gray matter ribbon onto the individual brain surface and then onto template surface meshes (i.e. “dtseries.nii” file), followed by surface-based smoothing using a geodesic Gaussian algorithm. Further details on fMRI data acquisition, task design and preprocessing can be found in (38,40). The task fMRI database includes six cognitive domains, which are emotion, language, motor, relational, social, and working memory. In total, there are 21 different experimental conditions. We excluded the two gambling conditions in our analysis due to the short event design of the gambling trials (1.5s for button press, 1s for feedback and 1s for ITI). The detailed description of the task paradigms as well as the selected cognitive domains can be found in (17,38)

### Decoding brain activity using graph convolution

A brain graph provides a network representation of brain organization by associating nodes with brain regions and defining edges via anatomical or functional connections (41). We recently found that convolutional operations on brain graph can be used to encode the within-network interactions of task-evoked brain responses and to decode a large number of cognitive tasks (17). Here, we proposed a more generalized form of graph convolution by using Chebyshev polynomials and explored how functional segregation and high-order functional interactions affects brain decoding.

### Step 1: Construction of brain graph

The decoding pipeline started with a weighted graph 𝒢 = (𝒱, ℰ, *W*), where 𝒱 is a parcellation of cerebral cortex into *N* regions, ℰ is a set of connections between each pair of brain regions, with its weights defined as *W*_*i,j*_. Many alternative approaches can be used to build such brain graph 𝒢, for instance using different brain parcellation schemes and constructing various types of brain connectomes (for a review, see (41)). Here, we used the Glasser’s multi-modal parcellation, consisting of 360 areas in the cerebral cortex, bounded by sharp changes in cortical architecture, function, connectivity, and topography (27). The edges between each pair of nodes were estimated by calculating the group averaged resting-state functional connectivity (RSFC) based on minimal preprocessed resting-state fMRI data from *N* = 1080 HCP subjects (Glasser et al., 2013). Additional preprocessing steps were applied before the calculation of RSFC, including regressing out the signals from white matter and csf, and bandpass temporal filtering on frequencies between 0.01 to 0.1 HZ. Functional connectivity was calculated on individual brains using Pearson correlation and then normalized using Fisher z-transform before averaging among the entire group of subjects. After that, a k-nearest-neighbor (k-NN) graph was built by only connecting each node to its 8 neighbors with the highest connectivity strength.

### Step 2: Mapping of brain activity onto the graph

After the construction of brain graph (i.e. defining brain parcels and edges), for each functional run and each subject, the preprocessed task-fMRI data was then mapped onto the set of brain parcels, resulting in a 2-dimensional time-series matrix. This time-series matrix was first split into multiple task blocks according to fMRI paradigms and then cut into sets of time-series of the chosen window size (e.g. 10 second). Shorter time windows were discarded in the process. The remaining time-series were treated as independent data samples during model training. As a result, we generated a large number of fMRI time-series matrices from all cognitive domains, i.e. a short time-series with duration of *T* for each of *N* brain parcels, *x* ∈ ℝ^*N*×*T*^. The entire dataset consists of over 1000 subjects for each cognitive domain (see Table S2 for detailed information), in total of 14,895 functional runs across the six cognitive domains, and 138,662 data samples of fMRI signals *x* ∈ ℝ^*N*×*T*^ when using a 10s time window (i.e. 15 functional volumes at TR=0.72s).

### Step 3: Spatiotemporal graph convolutions using ChebNet

Graph convolution relies on the graph Laplacian, which is a smooth operator characterizing the magnitude of signal changes between adjacent nodes. The normalized graph Laplacian is defined as:

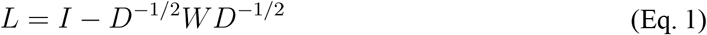

where D is a diagonal matrix of node degrees, I is the identity matrix, and W is the weight matrix. The eigendecomposition of Lapalcian matrix is defined as *L* = UΔU^*T*^, where U = (*u*_0_, *u*_1_, ⋯*u*_*N*−1_) is the matrix of Laplacian eigenvectors and is also called graph Fourier modes, and Δ = diag(*λ*_0_, *λ*_1_, ⋯*λ*_*N*−1_) is a diagonal matrix of the corresponding eigenvalues, specifying the frequency of the graph modes. In other words, the eigenvalues quantify the smoothness of signal changes on the graph, while the eigenvectors indicate the patterns of signal distribution on the graph.

For a signal *x* defined on graph, i.e. assigning a feature vector to each brain region, the convolution between the graph signal *x* ∈ ℝ^*N*×*T*^ and a graph filter *g*_*θ*_ ∈ ℝ^*N*×*T*^ based on graph 𝒢, is defined as their element-wise Hadamard product in the spectral domain, i.e.:

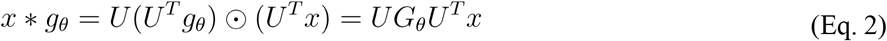

where *G*_*θ*_ = *diag*(*U*^*T*^*g*_*θ*_) and *θ* indicate a parametric model for graph convolution g_*θ*_, U = (*u*_0_, *u*_1_, ⋯*u*_*N*−1_) is the matrix of Laplacian eigenvectors and *U*^*T*^*x* is actually projecting the graph signal onto the full spectrum of graph modes. To avoid calculating the spectral decomposition of the graph Laplacian, ChebNet convolution (Defferrard et al., 2016) uses a truncated expansion of the Chebychev polynomials, which are defined recursively by:

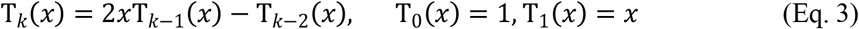

Consequently, the ChebNet graph convolution is defined as:

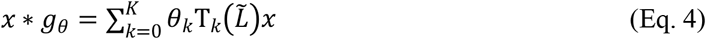

where 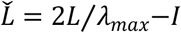 is a normalized version of graph Laplacian with *λ*_*max*_ being the largest eigenvalue, θ_k_ is the model parameter to be learned at each order of the Chebychev polynomials. It has been proved that the ChebNet graph convolution was naturally *K*-localized in space by taking up to *K* th order Chebychev polynomials (18), which means that each ChebNet convolutional layer integrates the context of brain activity within a *K*-step neighborhood.

### Brain-decoding pipeline using ChebNet graph convolution

We used a similar decoding pipeline as proposed in (17), consisting of 6 graph convolutional layers with 32 graph filters at each layer, followed by a flatten layer and 2 fully connected layers (256, 64 units). The model takes in a short series of fMRI volumes as input, maps the fMRI data onto the predefined brain graph and results in a 2-dimensional time-series matrix *X*^1^ ∈ ℝ^*N*×*T*^, i.e. a short time-series with duration of T for each of *N* brain parcels at the first ChebNet layer. The first ChebNet layer learns various shapes of temporal convolution kernels by treating multiple time steps as input channels (C = T) and propagates such temporal dynamics within (K=1) and between (K>1) brain networks. As a result, a set of “brain activation” maps are generated (see Figure 3-S1) and passed onto the next ChebNet layer for higher-order information integration (see Figure 3-S2), and so on. The learned graph representations in the last ChebNet layer were then imported to a 2-layer multilayer perceptron (MLP) to predict the cognitive state.

The entire dataset was split into training (60%), validation (20%), test (20%) sets using a subject-specific split scheme, which ensures that all fMRI data from the same subject was assigned to only one of the three sets. Approximately, the training set includes fMRI data from 700 unique subjects (depending on data availability for different cognitive tasks ranging from 1043 to 1085 subjects, see Table S2), with 176 subjects for validation set and 219 subjects for test set. Specifically, we used the training set to train model parameters, the validation set to evaluate the model at the end of each training epoch, and saved the best model with the highest prediction score on the validation set after 100 training epochs. The saved model was evaluated on the test set and reported the final decoding performance. We used Adam as the optimizer with the initial learning rate as 0.0001 on all cognitive domains. Additional l2 regularization of 0.0005 on weights was used to control model overfitting and the noise effect of fMRI signals. Dropout of 0.5 was additionally applied to the fully connected layers. The implementation of the ChebNet graph convolution was based on PyTorch 1.1.0, and was made publicly available in the following repository: https://github.com/zhangyu2ustc/gcn_tutorial_test.git.

### Effects of *K*-order in ChebNet brain decoding

ChebNet graph convolution used truncated expansion of Chebychev polynomials of order K for the approximation of graph convolution in the spectral space (see Eq. 4). The choice of K-order controls the scale of the information integration on the graph. When *K* = 0, *x* ∗ *g*_*θ*_ = *θ*_0_*x*, which indicates a global scaling factor on the input signal by treating each node independently, similar to the classical mass univariate analysis for brain activation detection. When *K* = 1, 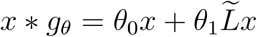, which indicates information integration between the direct neighbors and the current node on the graph (integrating signals within the same network). When *K* = 2, 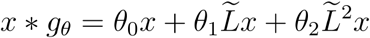, which indicates information integration within a two-step neighborhood on the graph (consisting of information from local area, within network and between networks). Generally speaking, when *K* > 1, the graph convolution integrates the information within a *K*-step neighbourhood by propagating graph signals not only within the same network but also among inter-connected brain networks. Thus, the *K*-order controls the propagation rate of information flow on the brain graph. We explored different choices of *K*-order in ChebNet spanning over the list of [0,1,2,5,10] and found a significant boost in both brain decoding and representational learning by using high-order graph convolutions.

### Saliency map of the decoding model: contribution of brain regions

The saliency map analysis aims to locate which part of the brain (or input features) helps to differentiate between cognitive tasks. We used a gradient approach named Guided BackProp (20) to visualize the contribution of inputs. This approach has been commonly used to visualize a deep neural network and easily generalized to graph convolutions. The basic idea is that if an input is relevant, a little variation on it will cause high change in the layer activation. This can be characterized by the gradient of the output given the input, with the positive gradients indicating that a small change to the input signals increases the output value. Specifically, for the graph signal *X*^1^ of layer and its gradient *R*^*l*^, the overwritten gradient 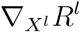 can be calculated as follows:

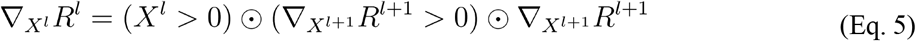

In order to generate the saliency map, we started from the output layer of a pre-trained model and used the above chain rule to propagate the gradients at each layer until reaching the input layer. This guided-backpropagation approach can provide a high-resolution saliency map which has the same dimension as the input data. We further calculated a heatmap of saliency by taking the variance across all time steps for each parcel, considering that the variance of the saliency curve provides a simplified way to evaluate the contribution of task-evoked hemodynamic response. To make it comparable across subjects, the saliency value was additionally normalized to the range [0,1], with its highest value at 1 (a dominant effect for task prediction) and lowest at 0 (no contribution to task prediction).

### Similarity analysis of layer representations in graph convolutions

ChebNet graph convolution maps the spatiotemporal dynamics of fMRI brain activity onto a new representational space in the spectral domain. Different representations were learned at each ChebNet graph convolutional layer by integrating the information flow within (*K* = 1) and between networks (*K* > 1). Besides, by using a multi-layer architecture, the scale of information integration was gradually enhanced, ranging from a local area (first ChebNet layer) to the whole-brain (last ChebNet layer). For a better understanding of the ChebNet models, we analyzed the similarity of learned representations between ChebNet layers as well as across different decoding models (e.g. using different K-orders). Considering the high-dimensional nature of learned representations (360*32 in our case), we evaluated the cross-layer and cross-model similarity using centered kernel alignment (CKA) with a linear kernel, which was proposed to compare layer representations of deep neural networks, not only in the same network trained from different initializations, but also across different models (21). Linear CKA is closely related to CCA and linear regression. Studies showed that CKA was invariant to orthogonal transformation and isotropic scaling, and consistently identified the correspondences of representations between layers, and thus can reveal pathology in neural networks representations (21). Here, we used CKA to evaluate the effects of K-orders in ChebNet brain decoding for both Motor and Working-memory tasks. First, we extracted the layer representations from each ChebNet layer by passing all data samples from the test set into the pre-trained decoding model and reshaped the representations (samples * nodes * channels) into a matrix *X* ∈ ℝ^*sample s*× *features*^. Then, the linear CKA of two representation matrices X and Y, either from different layers or different models, was defined as:

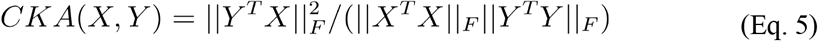

where 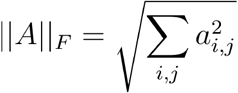 indicates the Frobenius norm of the cross-correlation matrix A. The CKA value was within the range [0,1], with its highest value at 1 (the same layer representation) and lowest at 0 (totally different layer representations).

### Projections of layer representations using t-SNE

For visualization purposes, we projected the high-dimensional layer representations to a 2-dimensional (2D) space by using t-SNE (25). Based on the t-SNE projections, we calculated the modularity score among task conditions as a measure of task segregation, representing the cost of brain state transition between task conditions. It has been shown that the modularity score on the shape graph constructed from individual fMRI data was significantly associated with participants’ in-scanner task performance (26). Here, we estimated the modularity score on t-SNE projections derived from not only fMRI signals but also layer representations of graph convolutional networks. Specifically, fMRI signals and layer representations were first mapped onto a 2D space by using t-SNE. Then, a k-NN graph (k=5) was constructed based on the coordinates of t-SNE projections by connecting each data sample with its five nearest neighbors in the 2D space. After that, a segregation index was defined by calculating the modularity score (Q) based on the partition of communities using task conditions, with a high separation value indicating more edges within the same task condition that expected by chance (42).

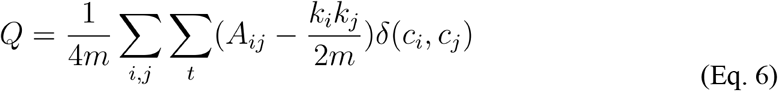

where *k*_*i*_ and *k*_*j*_ are the degrees of the nodes on the kNN graph, 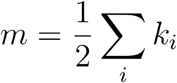 is the total number of edges on the kNN graph, and *δ*(*c*_*i*_, *c*_*j*_) indicates whether node *i* and node *j* belong to the same community (or task condition). The value of the task segregation index was within the range [-0.5,1], with the value close to 1 indicating a strong community structure and a positive value indicating the number of edges within the same task conditions exceeds the number expected by chance (e.g. on a random graph). The same procedure could also be applied to individual subject, i.e. constructing a kNN graph using t-SNE projections of fMRI signals or layer representations from the same subject. Their association with participants’ in-scanner task performance were also investigated by calculating the Pearson correlation coefficient of individual segregation index (i.e. modularity score on fMRI signals or layer representations) with the correct response and reaction time during working-memory tasks.

### Heritability analysis of brain representations and behavioral performances

For the heritability estimates of brain response and behavioral performance during WM tasks, we used the Sequential Oligogenic Linkage Analysis Routines (SOLAR) Eclipse software package (http://www.nitrc.org/projects/se_linux). SOLAR relies on the maximum variance decomposition of the covariance matrix Ω for a pedigree:

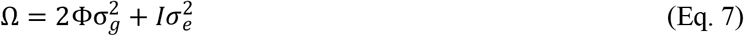

where 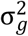 is the genetic variance due to the additive genetic factors, Φ is the kinship matrix representing the pairwise kinship coefficients among all individuals, 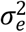 is the variance due to individual-specific environmental effects and measurement error, and *l* is an identity matrix. Narrow sense heritability is defined as the fraction of phenotypic variance 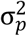 attributable to additive genetic factors: 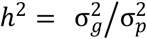. The significance of the heritability estimate is tested by comparing the likelihood of the model in which 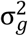 is constrained to zero with that of a model in which 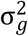 is estimated. Prior to the heritability estimation, all phenotype (brain and behavioral phenotypes) were adjusted for covariates including age, gender, handedness and head motion.

The heritability estimate was applied on 1070 subjects from HCP S1200 release with available behavioral and imaging data for WM tasks, which consist of 447 unique families, including 143 monozygotic-twin pairs, 83 dizygotic-twin pairs and 290 non-twin siblings.

We further performed the bivariate genetic analyses to quantify the shared genetic variance and the phenotypic correlation between brain responses and behavioral measures, relying on the following model:

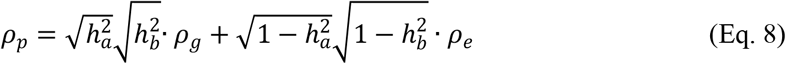

where Pearson’s phenotypic correlation *ρ*_*p*_ is decomposed into *ρ*_g_ and *ρ*_*e*_, where *ρ*_*g*_ is the proportion of variability due to shared genetic effects and *ρ*_*e*_ is that due to the environment, while 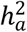 and 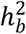 correspond to the narrow sense heritability for phenotypes a (representation of brain response) and b (behavioral scores).

## Acknowledgment

This work was partially supported by the Major Scientific Project of Zhejiang Lab (No. 2020ND8AD01), Courtois foundation through the Courtois NeuroMod Project and the IVADO Postdoctoral Scholarships Program. PB is supported by a salary award of “Fonds de recherche du Québec -Santé”, chercheur boursier junior 2.

## Author contributions

Conceptualization: YZ, PB; Methodology: YZ, PB; Visualization: YZ, PB;

Investigation: YZ, NF, AD, PB;

Writing—original draft: YZ, NF, PB

Writing—review & editing: YZ, NF, AD, PB

## Data and materials availability

We used the block-design task-fMRI dataset from the Human Connectome Project S1200 release, downloaded from https://db.humanconnectome.org/data/projects/HCP_1200. In total, fMRI data from 1095 unique subjects under six different task domains and resting-state were used in this study. The minimal preprocessed fMRI data of the CIFTI format were used, which maps individual fMRI time-series onto the standard surface template with 32k vertices per hemisphere. Our decoding pipeline, as well as the optimized decoding models and the construction of brain graphs, were made publicly available in the following repository: https://github.com/zhangyu2ustc/gcn_tutorial_test.git

## Supplementary Materials for

## Supplementary Figures

**Figure 1-S1.**
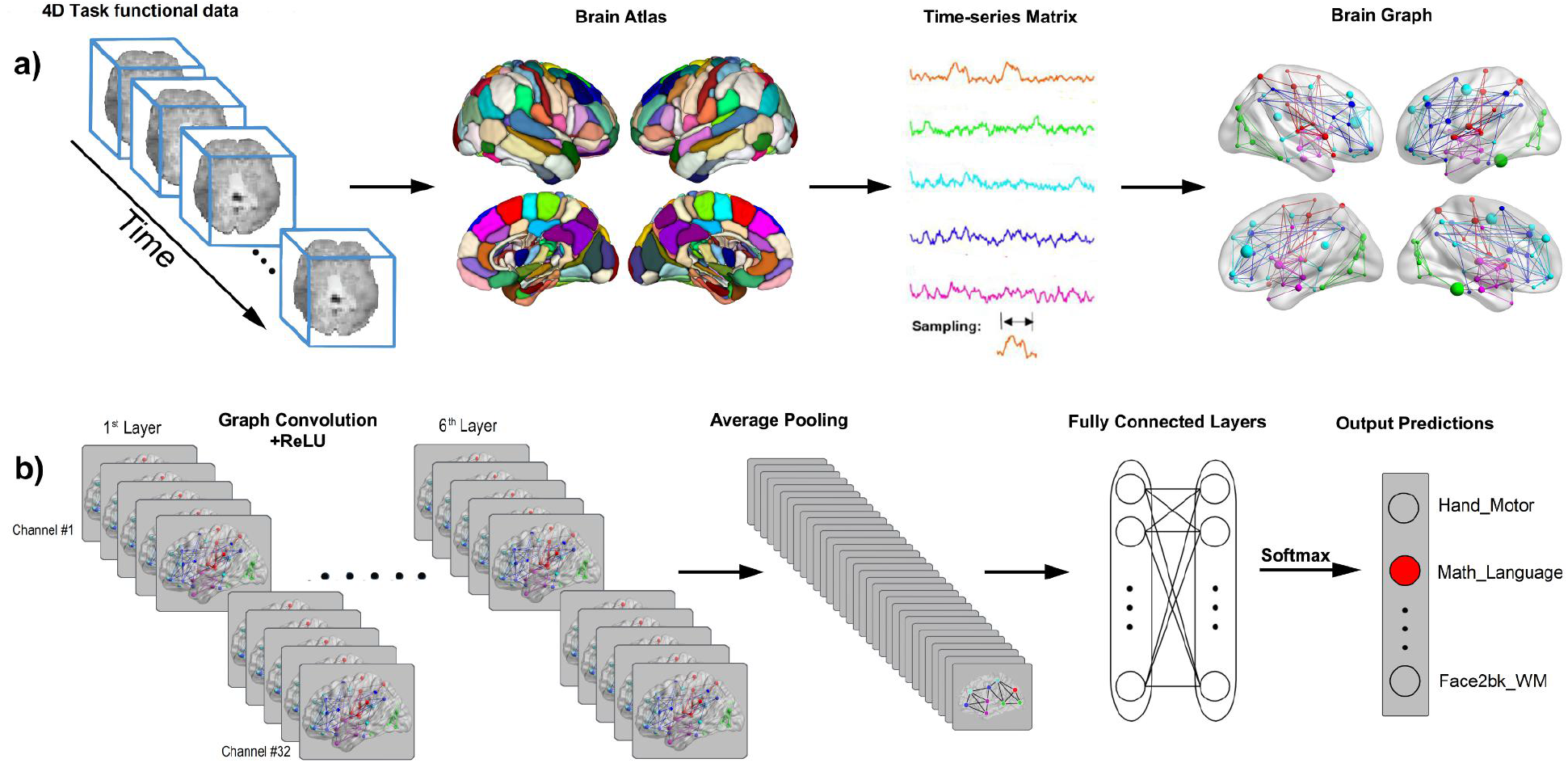
Pipeline of brain decoding using graph convolution network. The decoding model consists of six ChebNet graph convolutional layers with 32 graph filters at each layer, followed by a flatten layer and 2 fully connected layers. Specifically, for a short series of fMRI volumes, the measured brain activity was first mapped onto a predefined brain atlas consisting of hundreds of brain regions (e.g. 246 regions from Brainnetome atlas (43)). A functional graph was then constructed by calculating group-averaged resting-state functional connectivity for each pair of brain regions. Next, a new representation of task-evoked neural activity was generated through a multi-layer graph convolutional network, taking into account the segregation of localized brain activity and information integration among multiple brain networks. These representations were then used to predict the corresponding cognitive state associated with the short time window. The implementation of the ChebNet graph convolution was based on PyTorch 1.1.0, and was made publicly available in the following repository: https://github.com/zhangyu2ustc/gcn_tutorial_test.git.

**Figure 1-S2.**
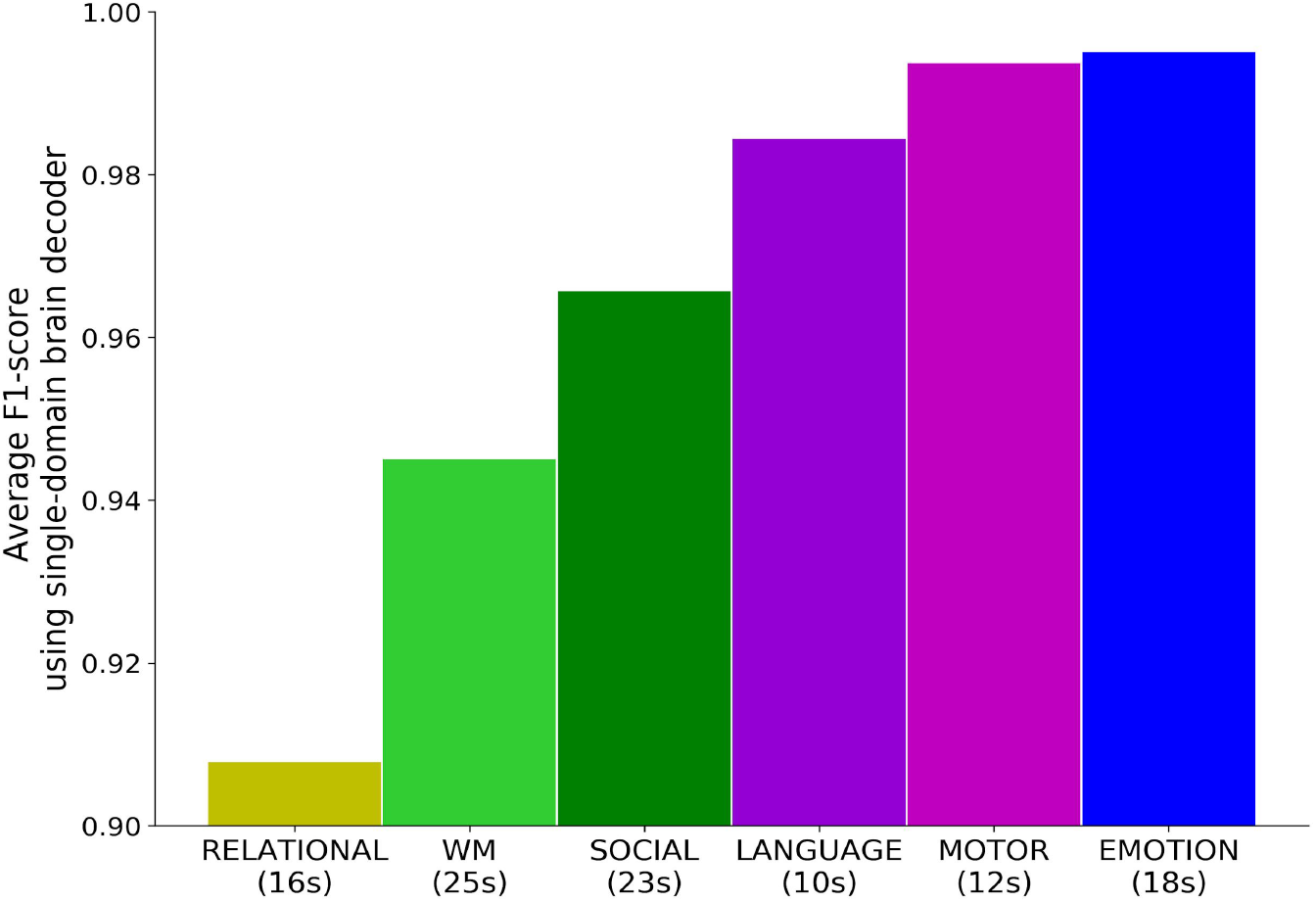
Decoding accuracy of single-domain brain decoders for each of the six cognitive domains. The same decoding pipeline was used as in Figure 1 except that the decoding model here was trained by using task-fMRI data exclusively from a single domain. Besides, variable temporal durations were used for each cognitive domain, according to the maximum length of event trials/blocks among all experimental tasks, for instance 12s for MOTOR tasks and 25s for WM tasks. Among the six cognitive domains, the emotion tasks (in blue, fearful face *vs* shape) and motor tasks (in magenta, distinguishing five types of body movements) were the most easily recognizable conditions, with F1-score reaching around 99%.

**Figure 3-S1.**
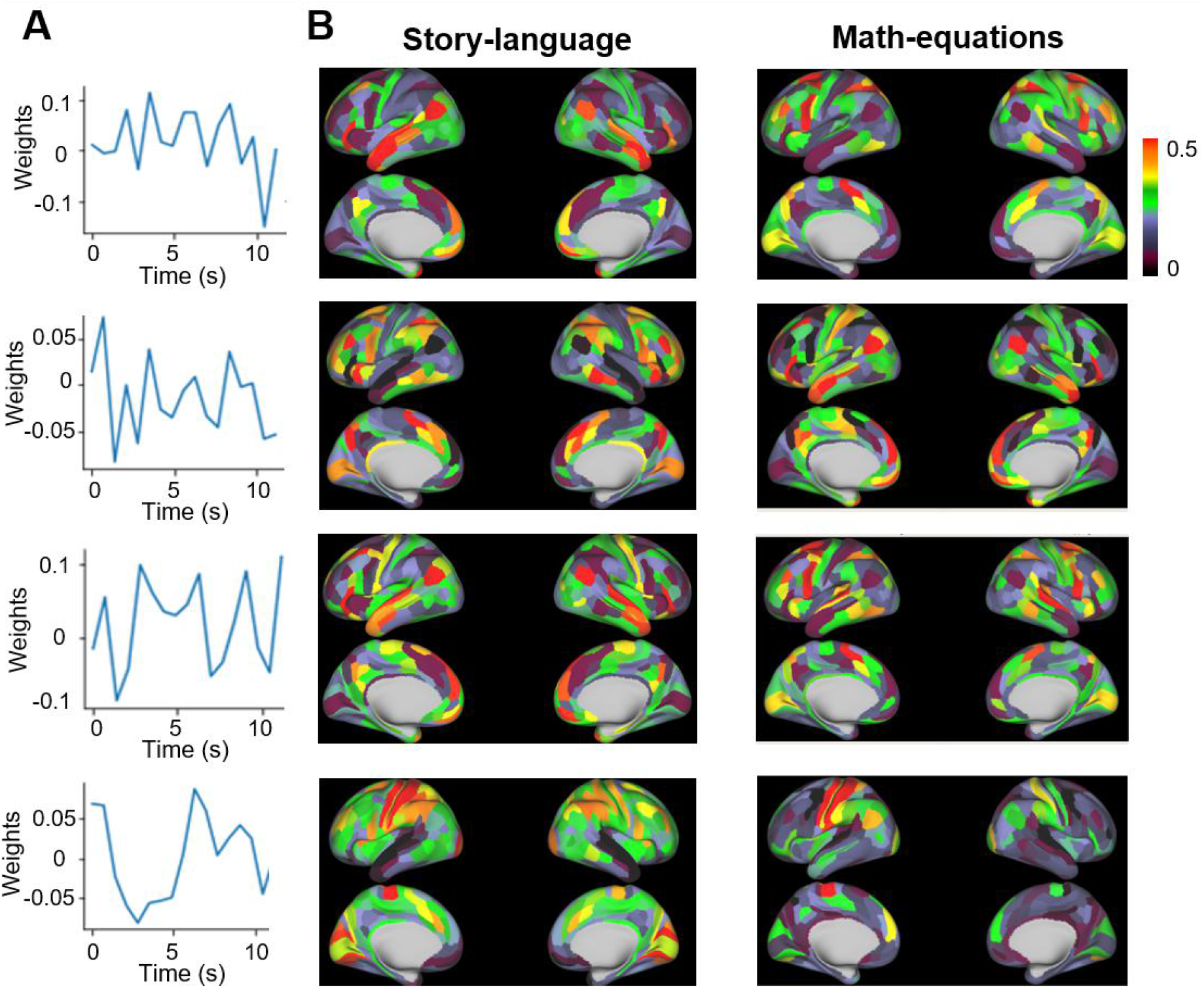
Temporal filters and the corresponding activation maps extracted from the first ChebNet layer for the Language tasks. We used the Language tasks as an example to illustrate that ChebNet graph convolutions captured hemodynamic response in the temporal domain and brain activations in the spatial domain. First of all, various temporal convolutional kernels were learned at the first ChebNet layer (1^st^ column), which resembled the hemodynamic response function in BOLD signals. Second, using these temporal filters, the corresponding “activation maps” in the first ChebNet layer were extracted and grouped according to the task conditions, e.g. story (2^nd^ column) and math (3^rd^ column) for the Language tasks. These activation maps demonstrated a possible explanation of the biological basis behind spatiotemporal graph convolutions. For instance, when the temporal filter only focused on brain activity within 0-6s after task onset (3^rd^ row), the classical language network and frontoparietal network were detected for the story and math condition respectively, responsible for language comprehension and numerical processing during the cognitive task. By contrast, when the temporal filter focused on brain activity 5-10s after task onset (4^th^ row), the motor and sensory cortex were detected for both conditions, responsible for button press during the task. When both time windows were taken into account (1^st^ row, 0-10s after task onset), the brain response in the language network and frontoparietal network were further enhanced while the activity in the auditory cortex was weakened. Similar analysis on the *Motor* tasks was shown in Figure 1-Supplement 2 in (17).

**Figure 3-S2.**
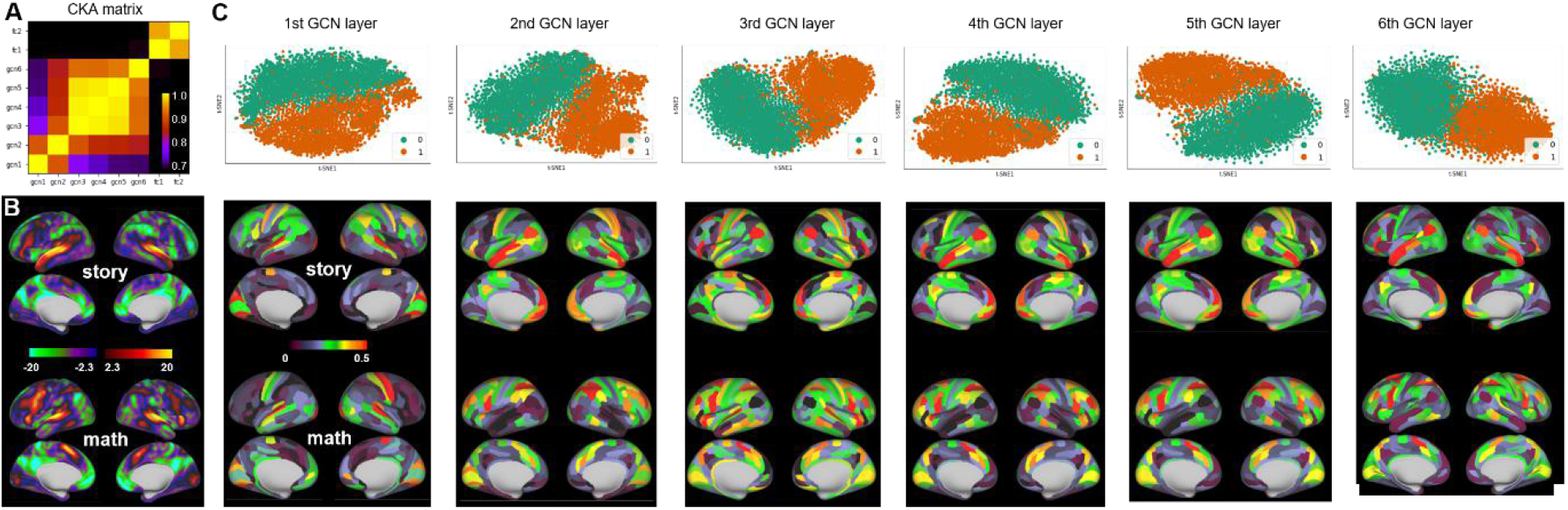
Layer representations in ChebNet resembled brain activation maps. We used the Language tasks as an example to illustrate the hierarchical organization of layer representations in ChebNet. At each ChebNet layer, the layer activations were extracted and saved as feature representations for the following analysis. First, the similarity of representations was calculated using CKA with a linear kernel (A), which illustrated a hierarchical organization of layer representations in ChebNet. These representations were then projected onto a 2-dimensional space using t-SNE (1^st^ row in C) which indicated a nice disassociation between different task conditions (e.g. story vs math for the Language task). After that, the “activation map” associated with each task condition (2^nd^ row in C) were calculated by averaging the layer representations across all data samples of the same category and mapped back onto the cortical surface. These representations resembled the actual brain activation maps detected by the canonical GLM approach (B), provided by (38), downloaded from neurovault (https://neurovault.org/collections/457/).

**Figure 4-S1.**
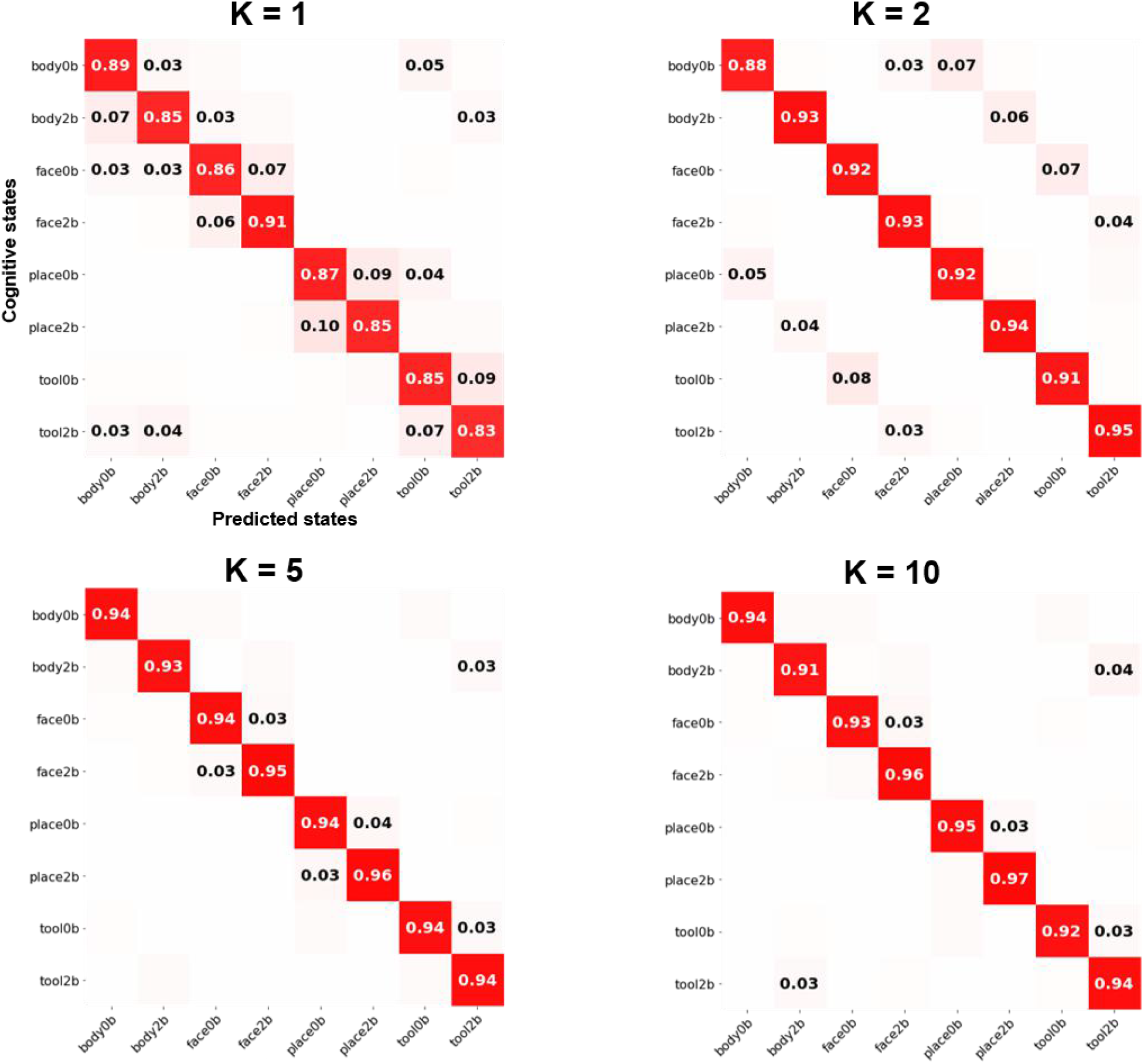
Confusion matrix of Working-memory tasks using ChebNet graph convolution with different K-orders. The confusion matrix was normalized by each task condition (row) such that each element in the matrix shows the recall score, i.e. among all predictions how many of them are positive predictions. The confusion matrix showed a nice block diagonal architecture, indicating that the majority of the cognitive tasks were accurately identified in all models. ALL decoding models were trained using 25s of fMRI time series. The classification errors were largely reduced when using high-order graph convolutions, e.g. K=1 vs K=5, especially for 0back vs 2back tasks. Our results indicated that large-scale functional integration of brain dynamics played an important role in the decoding of working memory tasks.

**Figure 6-S1.**
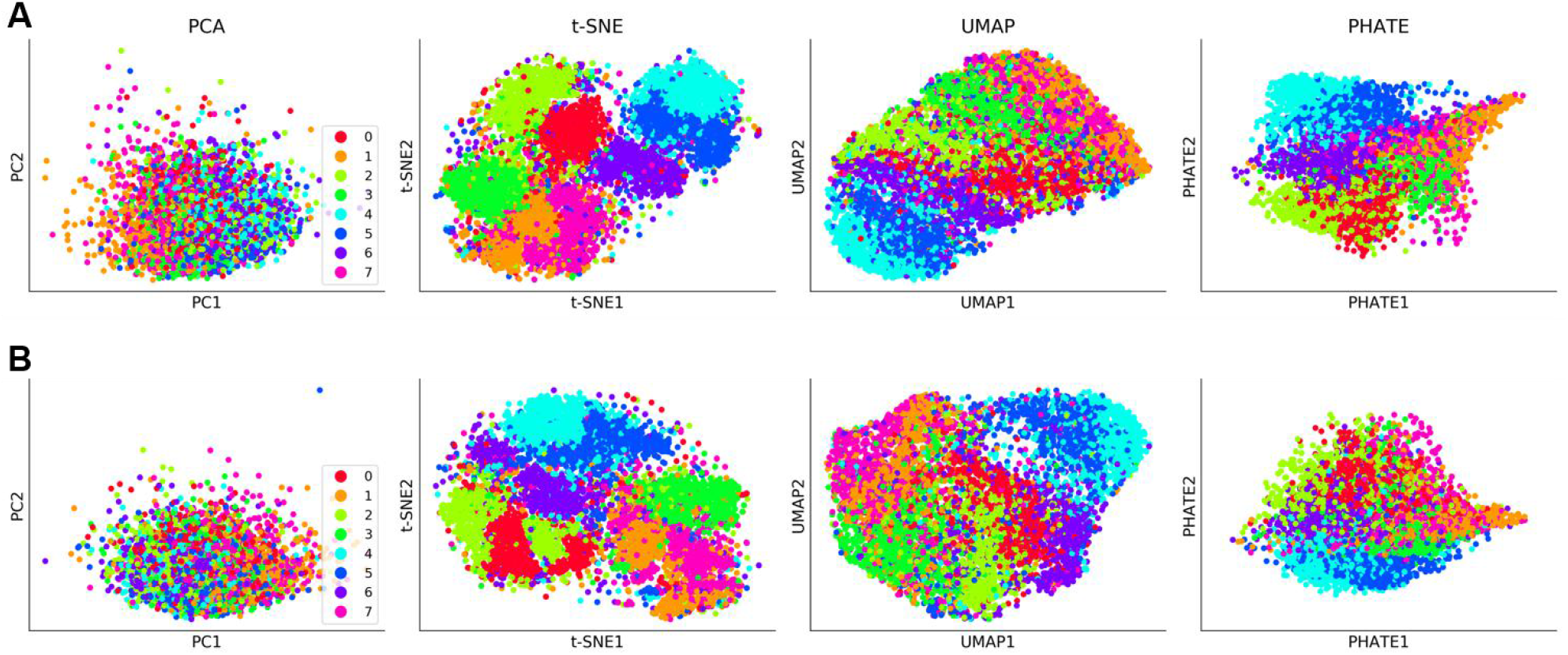
Projection of features representations from the ChebNet decoding model for WM tasks. Graph representations were mapped onto a 2-dimensional space by using different dimension reduction techniques, including PCA (first column), t-SNE (second column) (25), UMAP (third column) (44), and PHATE (last column)(45). The data samples included eight task conditions from Working-memory tasks, i.e. 0-back and 2-back on images of body parts (class 0 and 1), 0-back and 2-back on face images (class 2 and 3), 0-back and 2-back on place images (class 4 and 5), 0-back and 2-back on images of tools (class 6 and 7). Best visualization was provided by t-SNE. Two different decoding models were evaluated, including ChebNet-*K*5 (A) and ChebNet-*K*1 (B). The ChebNet-*K*5 model showed higher distinctions among eight WM task conditions.

**Figure 6-S2.**
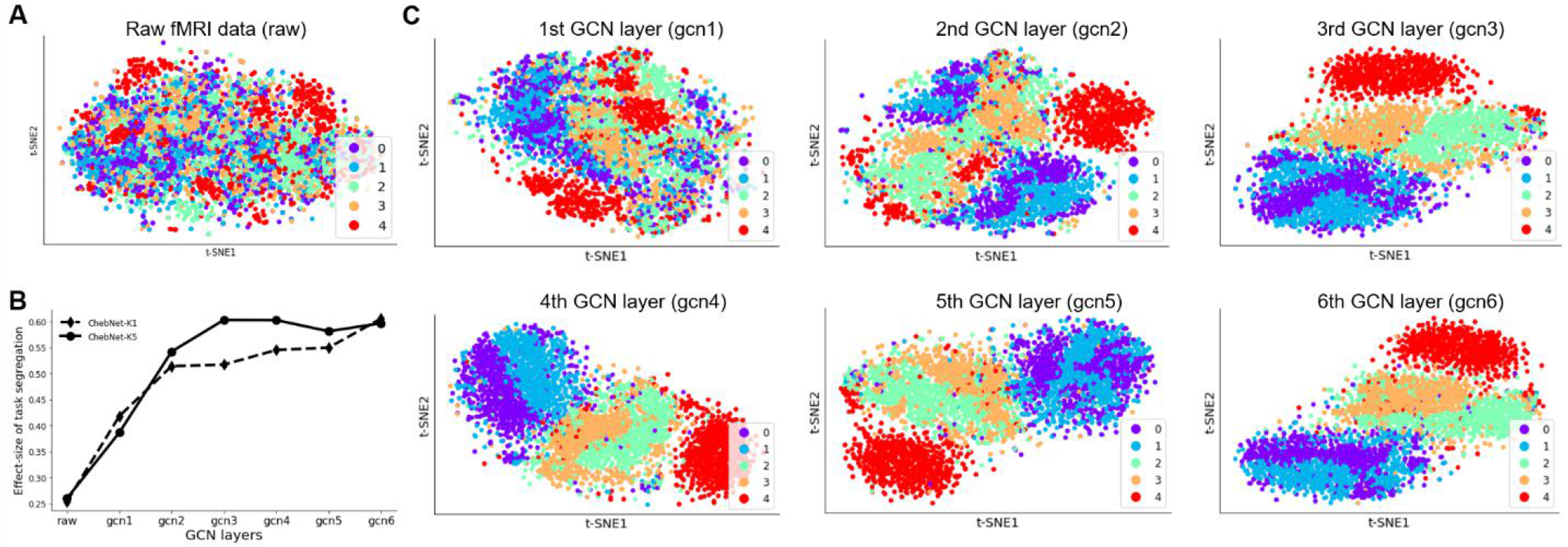
Layer representations of the ChebNet-*K*5 on the Motor task. Both raw fMRI data and layer representations of each ChebNet layer were projected onto 2-dimensional space by using t-SNE. No clear structure of task conditions was observed in the raw fMRI data (A), which slightly improved in the low-level representations, e.g. 1^st^ and 2^nd^ ChebNet layer (C). Since the 4th ChebNet layer, the tongue movement (in red) was easily distinguished from other motor tasks in the representations but still showed a mixture effect between left and right movements. In the last ChebNet layer, the representations for the five types of body movements were highly clustered and easily separated from each other (task segregation index Q =0.60). Similar level of task segregation in the last ChebNet layer when using different K-orders in ChebNet, but a faster convergence speed was detected in the ChebNet-*K*5 model (B). The Motor task data includes five types of body movements, i.e. the movement of right foot (class 0, in purple), left foot (class 1, in green), right hand (class 2, in cyan), left hand (class 3, in orange), and tongue (class 4, in red).

**Figure 7-S1.**
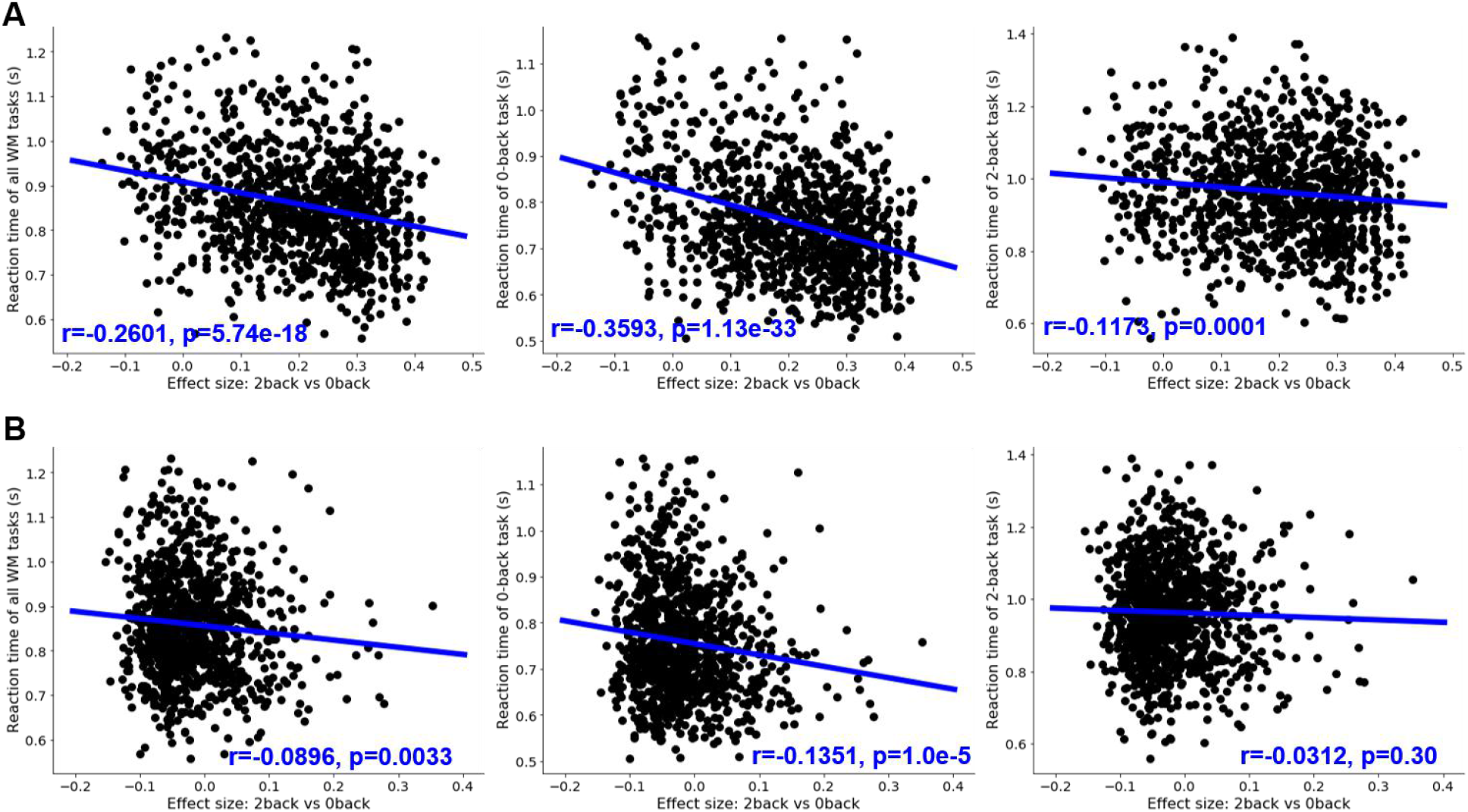
Modularity scores in the state-transition graph significantly correlated with the reaction time of Working-memory tasks. The modularity score was calculated based on the state-transition graph of each subject, as proposed by (26). We found significant correlations between the modularity scores of graph representations (A) and average reaction time during working-memory task (1^st^ panel), 0back task conditions (2^nd^ panel) and 2back task conditions (3^rd^ panel). Much weaker associations were detected in the raw fMRI data (B). The blue lines indicated the linear regression models between the modularity score of the state-transition graph and the reaction time during task performance. The analysis was done among all subjects from HCP S1200 release, with complete records of behavioral and imaging data for working memory tasks (N=1074).

## Supplementary Tables

**Table S1.** Scanning parameters and experimental designs of HCP task-fMRI dataset. The entire dataset includes in total of 14,895 functional runs across the six cognitive domains, and resulted in 138,662 data samples of fMRI signals when using a 10s time window (i.e. 15 functional volumes at TR=0.72s)

| **Task Domains** | **#Subjects** | **#Runs** | **#Volumes per run** | **#Trials per run** | **#Conditions** | **Minimal duration per block (sec)** |
| --- | --- | --- | --- | --- | --- | --- |
| Working memory | 1085 | 2 | 405 | 8 | 8 | 25 |
| Motor | 1083 | 2 | 284 | 10 | 5 | 12 |
| Language | 1051 | 2 | 316 | 8 | 2 | 10 |
| Social Cognition | 1051 | 2 | 274 | 5 | 2 | 23 |
| Relational processing | 1043 | 2 | 232 | 6 | 2 | 16 |
| Emotion | 1047 | 2 | 176 | 6 | 2 | 18 |

**Table S2.** Decoding accuracy from the single-domain decoders. Six single-domain decoders were trained by using fMRI responses from each cognitive domain exclusively and to predict the cognitive states on a trial basis. Different temporal durations were used, according to the maximum length of event trials on the target cognitive domain, for instance 12s for MOTOR tasks and 25s for WM tasks.

| **Task Domains** | **#Subjects** | **#Samples (single trials)** | **#Condition** | **Time windows (sec)** | **Decoding accuracy (F1-score)** |
| --- | --- | --- | --- | --- | --- |
| Working memory | 1085 | 17,360 | 8 | 25 | 0.9451 |
| Motor | 1083 | 21,660 | 5 | 12 | 0.9938 |
| Language | 1051 | 16,816 | 2 | 10 | 0.9845 |
| Social Cognition | 1051 | 10,510 | 2 | 23 | 0.9658 |
| Relational processing | 1043 | 12,516 | 2 | 16 | 0.9079 |
| Emotion | 1047 | 12,564 | 2 | 18 | 0.9952 |

**Table S3:** Comparison of decoding performance between different models. We reported the best performance for the baseline models after a grid search of the hyperparameters. For SVC approaches, we used the one-vs-rest (‘ovr’) decision function to handle multi-classes and reported the highest accuracy after the grid search for the hyper-parameter (C = [0.0001,0.001,0.1,1,10,100]). For Random Forest, we reported the highest accuracy after evaluating different settings of the classifier including depth of trees: [4,16,64,256,1024] and number of trees: [100,2000]. For MLP (multilayer perceptron), GCN (using first-order graph convolution, (17)) and ChebNet (using 5-order graph convolution), we reported the mean and standard deviation of the decoding accuracies among 10 fold cross-validation with shuffle splits. All models were evaluated on the task of decoding 21 task states by using 10s of fMRI signals (in total of 138,662 data samples).

| **Models** | **Train Accuracy** | **Validation Accuracy** | **Test Accuracy** |
| --- | --- | --- | --- |
| SVC-linear | 67.2% | 63.3% | 64.1% |
| SVC-rbf | 99.7% | 73.5% | 73.8% |
| Random Forest | 100% | 48.0% | 47.5% |
| MLP(256-64) | 87.9%(+/-1.83%) | 83.2%(+/-3.28%) | 76.1%(+/-0.41%) |
| GCN | 96.3%(+/-0.42%) | 90.2%(+/-0.21%) | 90.7%(+/-0.20%) |
| **ChebNet** | 90.91%(+/-0.18%) | **92.73(+/-0.12%)** | **93.43(+/-0.44%)** |

**Table S4:** Heritability analysis of brain responses and behavioral scores. Heritability estimates were conducted for both representations of brain responses and behavioral performance in-scanner, associated with WM tasks, after controlling for confounding effects of age, gender, handedness and head motion. The average accuracy (Acc) and reaction time (RT) showed high heritability estimates of additive genetic effects. For graph representations and raw fMRI signals, the high-dimensional data was first projected onto a 2-dimensional space using t-SNE and then the task segregation effect was estimated based on individual state-transition graph (see Method section). Significant heritability estimates were also detected in the ChebNet graph representations but not in raw fMRI signals.

| **Traits** | **$h^2$** | **SE** | **p-value** | **FDR corrected p-value** | **Covariance Explained** |
| --- | --- | --- | --- | --- | --- |
| **ChebNet graph representations** | **0.2882** | 0.0588 | **3.00E-07** | 3.43E-07 | 0.0017 |
| Raw fMRI signals | 0.0008 | 0.0545 | 0.4935 | 0.4935 | 0.0029 |
| WM_Task_Acc | 0.5624 | 0.0435 | 8.56E-27 | 3.42E-26 | 0.0434 |
| WM_Task_2bk_Acc | 0.5887 | 0.0425 | 6.97E-29 | 5.58E-28 | 0.0446 |
| WM_Task_0bk_Acc | 0.3215 | 0.0564 | 6.21E-09 | 8.29E-09 | 0.0222 |
| WM_Task_RT | 0.4118 | 0.0560 | 4.01E-13 | 8.01E-13 | 0.0105 |
| WM_Task_2bk_RT | 0.4534 | 0.0556 | 5.40E-15 | 1.44E-14 | 0.0094 |
| WM_Task_0bk_RT | 0.3294 | 0.0583 | 6.06E-09 | 8.29E-09 | 0.0085 |

## References

1. Haxby JV, Gobbini MI, Furey ML, Ishai A, Schouten JL, Pietrini P. Distributed and overlapping representations of faces and objects in ventral temporal cortex. Science. 2001 Sep 28;293(5539):2425–30.

2. Wang X, Liang X, Jiang Z, Nguchu BA, Zhou Y, Wang Y, et al. Decoding and mapping task states of the human brain via deep learning. Hum Brain Mapp. 2020 Apr 15;41(6):1505–19.

3. Li H, Fan Y. Interpretable, highly accurate brain decoding of subtly distinct brain states from functional MRI using intrinsic functional networks and long short-term memory recurrent neural networks. NeuroImage. 2019 Nov 15;202:116059.

4. Cohen JR, D’Esposito M. The Segregation and Integration of Distinct Brain Networks and Their Relationship to Cognition. J Neurosci. 2016 Nov 30;36(48):12083–94.

5. Deco G, Tononi G, Boly M, Kringelbach ML. Rethinking segregation and integration: contributions of whole-brain modelling. Nat Rev Neurosci. 2015 Jul;16(7):430–9.

6. Friston KJ. Functional and effective connectivity in neuroimaging: A synthesis. Hum Brain Mapp. 1994;2(1–2):56–78.

7. Tononi G, Sporns O, Edelman GM. A measure for brain complexity: relating functional segregation and integration in the nervous system. Proc Natl Acad Sci. 1994 May 24;91(11):5033–7.

8. Mottolese C, Richard N, Harquel S, Szathmari A, Sirigu A, Desmurget M. Mapping motor representations in the human cerebellum. Brain. 2013 Jan 1;136(1):330–42.

9. Haxby JV, Connolly AC, Guntupalli JS. Decoding Neural Representational Spaces Using Multivariate Pattern Analysis. Annu Rev Neurosci. 2014 Jul 8;37(1):435–56.

10. Eriksson J, Vogel EK, Lansner A, Bergström F, Nyberg L. Neurocognitive Architecture of Working Memory. Neuron. 2015 Oct 7;88(1):33–46.

11. Harrison SA, Tong F. Decoding reveals the contents of visual working memory in early visual areas. Nature. 2009 Apr;458(7238):632–5.

12. Riggall AC, Postle BR. The Relationship between Working Memory Storage and Elevated Activity as Measured with Functional Magnetic Resonance Imaging. J Neurosci. 2012 Sep 19;32(38):12990–8.

13. Christophel TB, Hebart MN, Haynes J-D. Decoding the Contents of Visual Short-Term Memory from Human Visual and Parietal Cortex. J Neurosci. 2012 Sep 19;32(38):12983–9.

14. Sligte IG, van Moorselaar D, Vandenbroucke ARE. Decoding the Contents of Visual Working Memory: Evidence for Process-Based and Content-Based Working Memory Areas? J Neurosci. 2013 Jan 23;33(4):1293–4.

15. Poldrack RA, Mumford JA, Schonberg T, Kalar D, Barman B, Yarkoni T. Discovering Relations Between Mind, Brain, and Mental Disorders Using Topic Mapping. Sporns O, editor. PLoS Comput Biol. 2012 Oct 11;8(10):e1002707.

16. Poldrack RA, Halchenko Y, Hanson SJ. Decoding the Large-Scale Structure of Brain Function by Classifying Mental States Across Individuals. Psychol Sci. 2009 Nov;20(11):1364–72.

17. Zhang Y, Tetrel L, Thirion B, Bellec P. Functional annotation of human cognitive states using deep graph convolution. NeuroImage. 2021 May 1;231:117847.

18. Defferrard M, Bresson X, Vandergheynst P. Convolutional Neural Networks on Graphs with Fast Localized Spectral Filtering. Adv Neural Inf Process Syst 29 [Internet]. 2016; Available from: http://arxiv.org/abs/1606.09375

19. Van Essen DC, Smith SM, Barch DM, Behrens TEJ, Yacoub E, Ugurbil K. The WU-Minn Human Connectome Project: An overview. NeuroImage. 2013 Oct 15;80:62–79.

20. Springenberg JT, Dosovitskiy A, Brox T, Riedmiller M. Striving for Simplicity: The All Convolutional Net. 2014 Dec 21 [cited 2021 Jan 27]; Available from: https://arxiv.org/abs/1412.6806v3

21. Kornblith S, Norouzi M, Lee H, Hinton G. Similarity of Neural Network Representations Revisited. ArXiv190500414 Cs Q-Bio Stat [Internet]. 2019 Jul 19 [cited 2021 Jan 15]; Available from: http://arxiv.org/abs/1905.00414

22. Nie Q-Y, Müller HJ, Conci M. Hierarchical organization in visual working memory: From global ensemble to individual object structure. Cognition. 2017 Feb 1;159:85–96.

23. Raut RV, Snyder AZ, Raichle ME. Hierarchical dynamics as a macroscopic organizing principle of the human brain. Proc Natl Acad Sci. 2020 Aug 25;117(34):20890–7.

24. Bressler SL, Menon V. Large-scale brain networks in cognition: emerging methods and principles. Trends Cogn Sci. 2010 Jun 1;14(6):277–90.

25. Maaten L van der, Hinton G. Visualizing Data using t-SNE. J Mach Learn Res. 2008;9(86):2579–605.

26. Saggar M, Sporns O, Gonzalez-Castillo J, Bandettini PA, Carlsson G, Glover G, et al. Towards a new approach to reveal dynamical organization of the brain using topological data analysis. Nat Commun. 2018 Dec;9(1):1399.

27. Glasser MF, Coalson TS, Robinson EC, Hacker CD, Harwell J, Yacoub E, et al. A multi-modal parcellation of human cerebral cortex. Nature. 2016 Aug;536(7615):171–8.

28. Haxby JV, Guntupalli JS, Connolly AC, Halchenko YO, Conroy BR, Gobbini MI, et al. A Common, High-Dimensional Model of the Representational Space in Human Ventral Temporal Cortex. Neuron. 2011 Oct;72(2):404–16.

29. Nishimoto S, Vu AT, Naselaris T, Benjamini Y, Yu B, Gallant JL. Reconstructing visual experiences from brain activity evoked by natural movies. Curr Biol. 2011 Oct 11;21(19):1641–6.

30. Mitchell TM, Shinkareva SV, Carlson A, Chang K-M, Malave VL, Mason RA, et al. Predicting Human Brain Activity Associated with the Meanings of Nouns. Science. 2008 May 30;320(5880):1191–5.

31. Huth AG, Nishimoto S, Vu AT, Gallant JL. A continuous semantic space describes the representation of thousands of object and action categories across the human brain. Neuron. 2012 Dec 20;76(6):1210–24.

32. Varoquaux G, Schwartz Y, Poldrack RA, Gauthier B, Bzdok D, Poline J-B, et al. Atlases of cognition with large-scale human brain mapping. Diedrichsen J, editor. PLOS Comput Biol. 2018 Nov 29;14(11):e1006565.

33. Yoshimura N, Tsuda H, Kawase T, Kambara H, Koike Y. Decoding finger movement in humans using synergy of EEG cortical current signals. Sci Rep. 2017 Sep 12;7(1):11382.

34. Poldrack RA. Inferring Mental States from Neuroimaging Data: From Reverse Inference to Large-Scale Decoding. Neuron. 2011 Dec;72(5):692–7.

35. Cole MW, Ito T, Bassett DS, Schultz DH. Activity flow over resting-state networks shapes cognitive task activations. Nat Neurosci. 2016;12.

36. Penfield W, Boldrey E. SOMATIC MOTOR AND SENSORY REPRESENTATION IN THE CEREBRAL CORTEX OF MAN AS STUDIED BY ELECTRICAL STIMULATION1. Brain. 1937 Dec 1;60(4):389–443.

37. Golarai G, Ghahremani DG, Whitfield-Gabrieli S, Reiss A, Eberhardt JL, Gabrieli JD, et al. Differential development of high-level visual cortex correlates with category-specific recognition memory. Nat Neurosci. 2007 Apr;10(4):512–22.

38. Barch DM, Burgess GC, Harms MP, Petersen SE, Schlaggar BL, Corbetta M, et al. Function in the human connectome: Task-fMRI and individual differences in behavior. NeuroImage. 2013 Oct 15;80:169–89.

39. Bijsterbosch J, Harrison SJ, Jbabdi S, Woolrich M, Beckmann C, Smith S, et al. Challenges and future directions for representations of functional brain organization. Nat Neurosci. 2020 Dec;23(12):1484–95.

40. Glasser MF, Sotiropoulos SN, Wilson JA, Coalson TS, Fischl B, Andersson JL, et al. The minimal preprocessing pipelines for the Human Connectome Project. NeuroImage. 2013 Oct 15;80:105–24.

41. Bullmore E, Sporns O. Complex brain networks: graph theoretical analysis of structural and functional systems. Nat Rev Neurosci. 2009 Mar;10(3):186–98.

42. Newman MEJ. Modularity and community structure in networks. Proc Natl Acad Sci. 2006 Jun 6;103(23):8577–82.

43. Fan L, Li H, Zhuo J, Zhang Y, Wang J, Chen L, et al. The Human Brainnetome Atlas: A New Brain Atlas Based on Connectional Architecture. Cereb Cortex. 2016 Jan 8;26(8):3508–26.

44. McInnes L, Healy J, Saul N, Großberger L. UMAP: Uniform Manifold Approximation and Projection. J Open Source Softw. 2018 Sep 2;3(29):861.

45. Moon KR, van Dijk D, Wang Z, Gigante S, Burkhardt DB, Chen WS, et al. Visualizing structure and transitions in high-dimensional biological data. Nat Biotechnol. 2019 Dec;37(12):1482–92.

